# Quantitative Spatial Analysis of Chromatin Biomolecular Condensates using Cryo-Electron Tomography

**DOI:** 10.1101/2024.12.01.626131

**Authors:** Huabin Zhou, Joshua Hutchings, Momoko Shiozaki, Xiaowei Zhao, Lynda K. Doolittle, Shixin Yang, Rui Yan, Nikki Jean, Margot Riggi, Zhiheng Yu, Elizabeth Villa, Michael K. Rosen

## Abstract

Phase separation is an important mechanism to generate certain biomolecular condensates and organize the cell interior. Condensate formation and function remain incompletely understood due to difficulties in visualizing the condensate interior at high resolution. Here we analyzed the structure of biochemically reconstituted chromatin condensates through cryo-electron tomography. We found that traditional blotting methods of sample preparation were inadequate, and high-pressure freezing plus focused ion beam milling was essential to maintain condensate integrity. To identify densely packed molecules within the condensate, we integrated deep learning-based segmentation with novel context-aware template matching. Our approaches were developed on chromatin condensates, and were also effective on condensed regions of in situ native chromatin. Using these methods, we determined the average structure of nucleosomes to 6.1 and 12 Å resolution in reconstituted and native systems, respectively, and found that nucleosomes form heterogeneous interaction networks in both cases. Our methods should be applicable to diverse biochemically reconstituted biomolecular condensates and to some condensates in cells.

## Introduction

Biomolecular condensates are increasingly recognized for their important roles in myriad biological processes, ranging from gene expression and signal transduction to stress responses (1–3). Dysregulation of condensates is implicated in diseases including neurodegenerative disorders (e.g. amyotrophic lateral sclerosis), cancer and viral infection (4–6). The internal structure and dynamic rearrangements of condensates are believed to play pivotal roles in the normal functions of condensates and in disease progression (1, 2).

Analogous to traditional structural analyses of molecular machines, direct visualization of components within condensates could provide substantial insights into the mechanisms by which condensates form, respond to signals, and function. This goal encompasses characterizing the conformations of individual molecules and their discrete complexes, and also understanding the higher-order organization of these components, encompassing their positions, orientations, and interactions. Notably, this latter goal is different from most NMR, crystallographic, and single-particle cryo-electron microscopy (cryo-EM) analyses, which generally seek to understand the average structure of individual components at high resolution rather than the spatial arrangements of molecules in the collection.

Biochemical reconstitution of condensates offers a controlled environment that simplifies these complex systems, allowing for detailed studies of their activities and the derivation of underlying principles applicable to more complex in vivo systems (7–10). Structural studies of such reconstituted systems are particularly promising as they provide a precise understanding of structure-function relationships in a well-defined context where all component parts are known a priori.

Cryo-electron tomography (cryo-ET) is a potentially powerful technique for structural analyses of condensates because it enables visualization of all individual particles within a field of view (11). Particles can be computationally combined to yield an average structure, and also their spatial arrangements can be analyzed to yield information on higher-order organization. However, unique and generic challenges arise when studying condensates with cryo-ET. Many condensates behave as dynamic liquids, in which molecules interact weakly and transiently (1–3). This property makes them particularly susceptible to perturbations during sample preparation, especially in biochemically reconstituted systems (12, 13).The high density of molecules within condensates can also obscure individual molecular details and produce overlapping electron density in low electron dosed, noisy tomograms. Furthermore, to achieve high resolution reconstructions of individual molecules/complexes, traditional single-particle cryo-EM approaches typically discard the vast majority of particles (14), and cryo-ET approaches compensate for missing wedge artifacts by averaging particles in different orientations (11, 15). However, these procedures are inappropriate when a key goal of a study is to identify all particles in a sample and understand their spatial organization. Because of these issues, while a number of studies of both biochemically reconstituted (16–19) and cellular (20–22) condensates have substantially benefitted from cryo-ET, these analyses have not yet yielded high resolution structures of condensate components or a quantitative understanding of the spatial relationships between molecules.

We sought to address these challenges through cryo-ET studies of condensates formed by polynucleosome arrays, which model cellular chromatin from the eukaryotic nucleus (13, 23, 24). We selected this condensate system for several reasons. First, chromatin condensates are representative of the many condensates that form through multiva-lency-driven liquid-liquid phase separation (LLPS) (23, 25). Second, they present significant technical advantages for cryo-ET, as individual nucleosomes are flattened discs of known structure, consisting of an ~120 kDa proteinaceous core wrapped twice by ~96 kDa (~147 base pairs) of double stranded DNA(26). Thus, nucleosomes are easily distinguished and identified within tomograms due to their large, unique electron-dense shape. Finally, both the structure of individual nucleosomes and their higher-order organization are important for numerous genomic functions (27–29), and chromatin condensates represent a powerful biomimetic system to understand how nuclear biochemistry is affected by the chromatin environment (23). Several studies have investigated the structure of native chromatin fibers using cryo-electron microscopy. However, most of these have concentrated on the conformations of individual nucleosomes or fibers rather than the higher order organization of these units or the network of interactions between them (17, 30–34).

Using chromatin condensates, we developed a pipeline spanning from sample preparation to image analysis that enables structural investigations of the condensate interior. The pipeline is effective for both biochemically reconstituted chromatin condensates and native chromatin in isolated mammalian cell nuclei and intact mammalian cells. Using it, we determined the average structure of nucleosomes to 6.1 and 12 Å resolution from the tomography data in reconstituted and native systems, respectively, and show that nucleosomes have a nearly random orientation distribution in both cases. Our methodology should be applicable to many reconstituted condensates and to certain cellular condensates where components are large and distinctive.

## Results

### Blotting and Self-wicking Distort Condensate Morphologies

To visualize condensates using cryo-ET, the sample must be sufficiently thin to allow electron transmission. Initially, we applied standard blotting techniques, where solutions containing chromatin condensates were pipetted onto grids, and excess liquid was removed with filter paper to create a thin layer (Fig 1a). Subsequent plunge freezing and tilt-series collection enabled reconstruction of tomograms with high contrast (Fig 1b). However, this process substantially distorted the chromatin condensate structure. The droplets were compressed into a thin layer, which, coupled with capillary forces, caused significant deformation (Fig 1b). The droplets were no longer round, contained numerous internal cavities, and consistently adhered to the carbon support on the grid. Extensive naked DNA was also observed, indicating that nucleosomes had been disassembled. These artifacts compromised the reliability of the structural data. Attempts to mitigate damage using a one-sided blotting technique were unsuccessful; droplets still adhered to the carbon in unusual shapes (Fig S1a, b).

**Figure 1.**
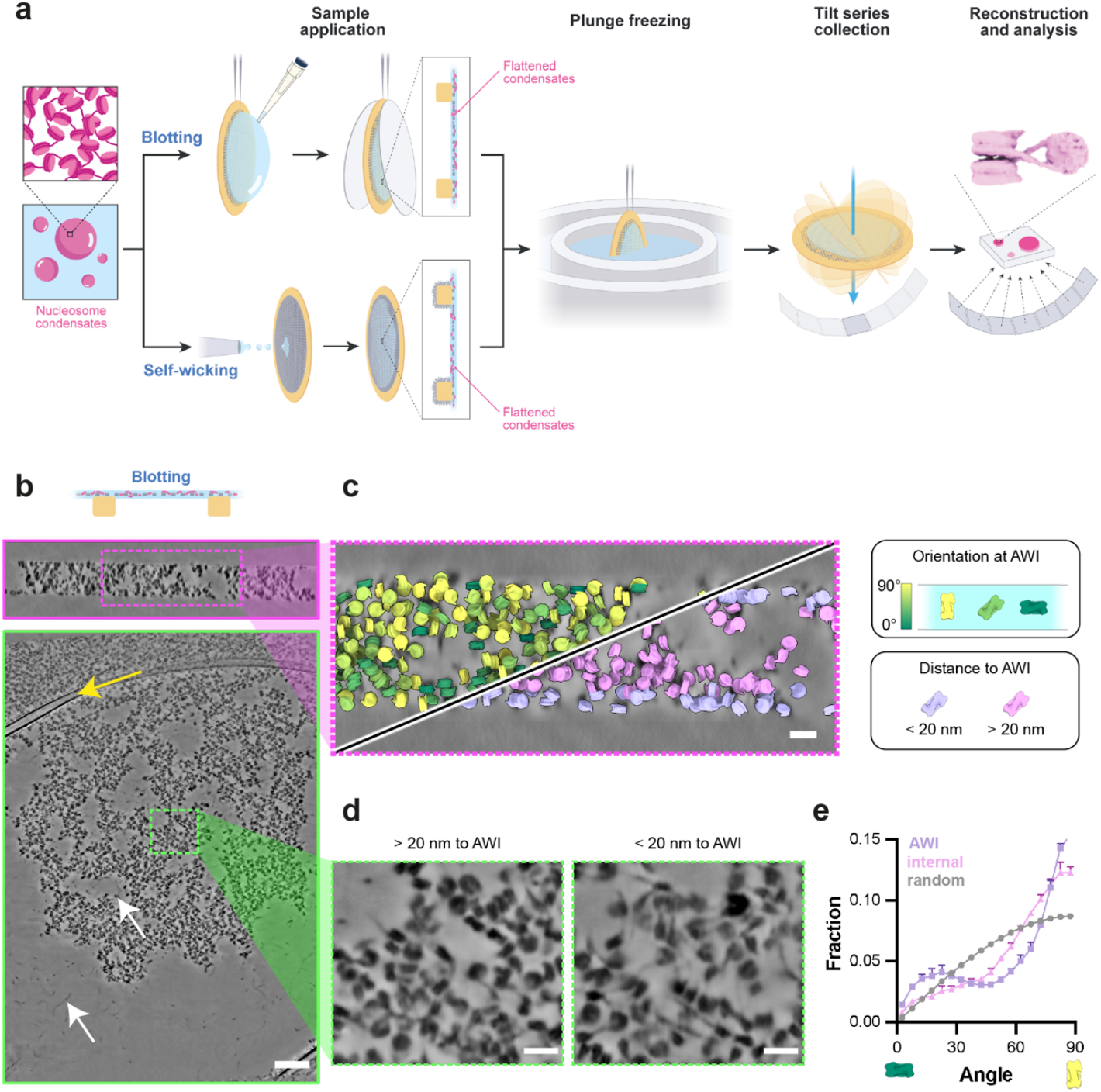
Morphological distortions in condensates by blotting and self-wicking techniques. **(a)** Diagram depicting blotting- and self-wicking-based sample preparation for cryo-ET. **(b)** Orthogonal cross-sections of chromatin condensates processed using the blotting technique. Yellow arrows indicate carbon support film on the grids. White arrows indicate exposed nucleosomes and bare DNA regions. Scale bar is100 nm. X-Y, X-Z views are shown with green, magenta outlines, respectively. **(c)** Zoom-in the X-Z view of the tomogram, with the assigned nucleosomes colored by the relative orientation (top left) or distance (bottom right) to the AWI. **(d)** Representative sections demonstrating chromatin condensates located close to and away from the air-water interface. Scale bars are 20 nm. **(e)** Angular distribution relative to beam direction (Z-axis) of nucleosomes at the air-water interface and in the core of the condensate.

Subsequently, we explored the self-wicking method (35), in which a small volume of sample is sprayed on the grid and nanofibers on the grid then extract excess buffer to reduce specimen thickness (Fig 1a). This approach presented similar artifacts; droplets continued to adhere to the carbon support, were compressed into thin layers, and exposed significant amounts of naked DNA (Fig S1c, d). As described below, blotting and self-wicking also altered the density of nucleosomes in the condensates.

An additional problem with both the blotting and self-wicking approaches is that condensates at the air-water interface (AWI) displayed a marked orientational bias. Context aware template matching (CATM; detailed below) analysis of nucleosome orientations revealed that nucleosomes throughout the sample had an orientation distribution that is highly non-random, contrary to expectations for an isotropic condensate (see below). Those at the AWI present disproportionately have their faces parallel to the interface (Fig 1c-e, S1d, e).

To further investigate this bias, we blotted non-phase separated samples composed of mononucleosomes and collected tomograms (Fig S2a). The resulting images confirmed that nucleosomes were also predominantly trapped at the AWI in a face-on orientation with unwrapped nucleosomes (Fig S2b, c). Preferred orientation due to the AWI is a well documented issue in single-particle cryo-EM (36); our results further highlight the challenges posed by these sample preparation methods in preserving native higher-order organization of biochemically reconstituted condensates.

### Preservation of Droplet Morphology via High-Pressure Freezing and Cryo-focused Ion Beam Milling

To best preserve condensates in their native state and during sample preparation, we turned to high-pressure freezing (HPF) and cryo-focused ion beam (Cryo-FIB) milling using the Waffle Method (37). Condensate-containing samples were transferred to the back side of holey carbon grids, and droplets (~0.5-3 µm in diameter) were allowed to settle on the carbon film for 30 seconds before freezing to enrich droplets on the front side of the grid. After HPF, such droplets were fully embedded in the 25 µm thick slab of vitreous ice on the grid (Fig 2a).

**Figure 2.**
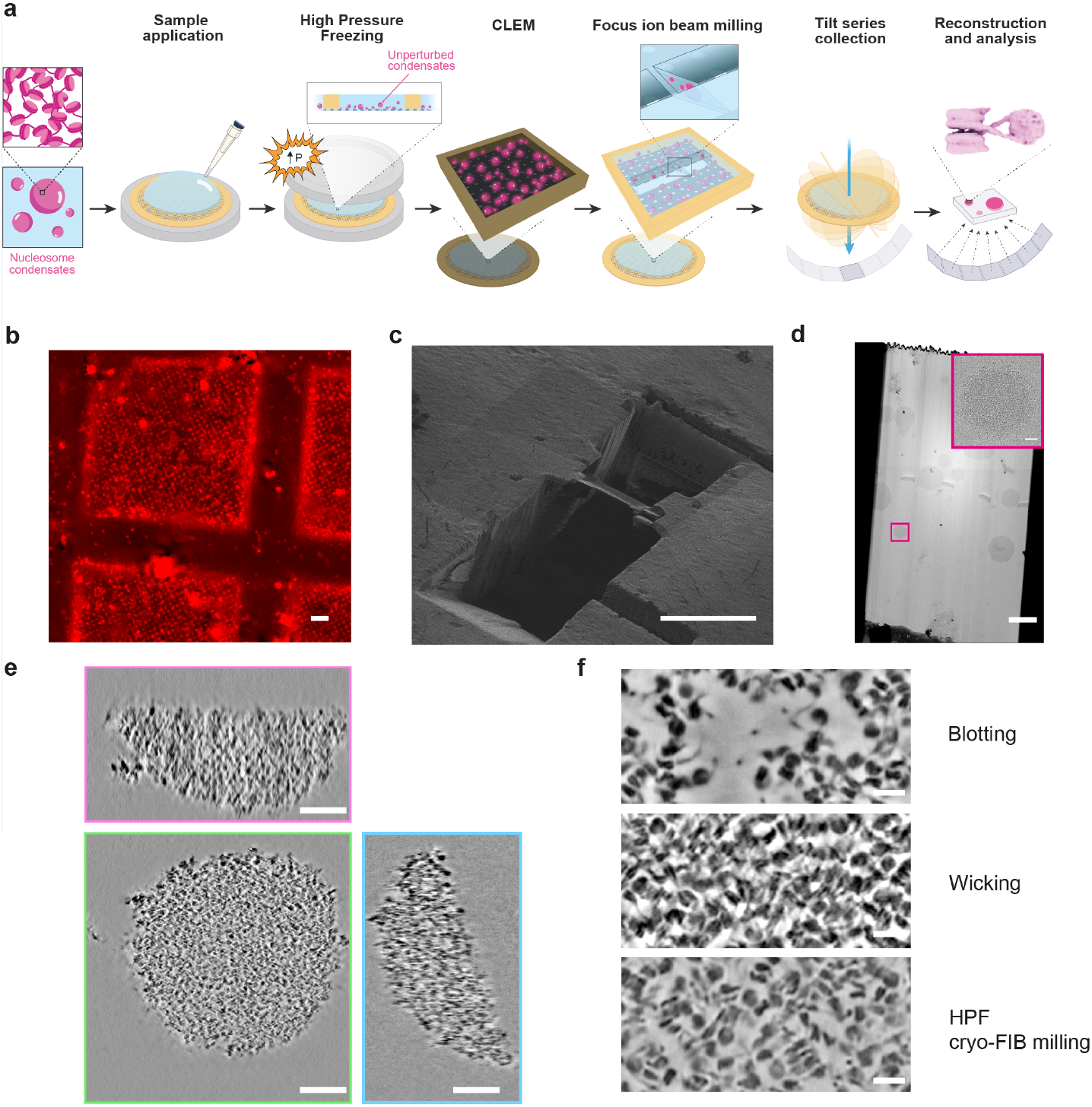
Preservation of droplet morphology via high-pressure freezing and cryo-focused ion beam milling. **(a)** Diagram depicting the integrated approach employing high-pressure freezing, correlative light and electron microscopy, and cryo-electron tomography to visualize chromatin droplets in their native state. Large condensates cut into discs, and small condensates remain fully round. **(b)** Representative squares of cryo-fluorescent images. Chromatin was labeled on histone H2B with AF594. Scale bar is 20 μm. **(c)** An example of the lamella production using FIB-milling. Scale bar is 20 μm. **(d)** Representative images of medium magnification montage for lamella using cryo-TEM, with a scale bar represent 2 μm. Insert in the enlarge image of the red box, the scale bar is 200 nm. **(e)** Orthogonal tomographic cross-sections of chromatin condensates prepared via HPF. Scale bar is 100 nm. X-Y, X-Z and Y-Z views are shown with green, red and blue outlines, respectively. **(f)** Tomographic sections illustrating chromatin in samples prepared using the indicated methods. Scale bar is 20 nm.

We then used cryo-fluorescence microscopy of the frozen grids with Alexa Fluor 594 (AF594)-labeled chromatin condensates to identify grid squares, and subregions within them, with highly abundant droplets in order to increase the probability of capturing condensates in the milled lamella (Fig 2b). The grids were transferred to the chamber of the Cryo-FIB instrument and the droplet-enriched regions were milled at random sub-locations (Fig 2c). Droplets within the lamella were identified via transmission electron microscopy (Fig. 2d). Effective template matching (see below) required lamella with 80-150 nm thickness. This thickness was important, since thicker samples resulted in increased electron scattering, leading to lower resolution and poor contrast, which compromised the quality of the acquired images and hindered accurate structural analysis (38). Using HPF and cryo-FIB milling (Movie 1), we could solve many of the problems associated with blotting and self-wicking methods. HPF produced spherical droplets with little apparent distortion (Fig 2e). These could be generated from condensates that were free-floating in solution and not attached to the carbon supports of the grid. Internally, nucleosomes were relatively uniform in density and the droplets lacked large cavities. No naked DNA could be observed, indicating that the nucleosomes were generally intact. Finally, the overall density of HPF condensates was intermediate between blotted samples, where disruption generally spread nucleosomes, and wicked samples, where dehydration increased nucleosome packing (Fig 2f).

### Assigning Nucleosomes in Synthetic Tomograms with Context-Aware Template Matching (CATM)

Understanding molecular interactions and higher order organization within condensates requires accurate assignments of molecular identity with high coverage in tomograms. This is a challenging task due to the crowded environment inside condensates and technical issues including the “missing wedge” problem and contrast transfer function (CTF) modulation, which render molecular identities ambiguous even for well-characterized structures (39) (Fig 3a, S3). To evaluate the efficacy of various template matching procedures in identifying individual nucleosomes in condensates, we first simulated tomograms with a mix of nucleosomes and DNA, mirroring the density of the chromatin condensates and the noise level (40) in our images (Fig S4a). Using standard template matching techniques (TM) (41, 42) (see Methods) applied to these simulated data (Fig 3c), we positioned the center of mass of the template at each voxel, rotated it at various angles, applied the missing wedge, and performed cross-correlation. The highest cross-correlation coefficient (CCC) values then determined the center of mass voxel and orientation of each matched template. In refinement, when two particles had centers of mass within a specified distance cutoff (4 nm, slightly smaller than the width of nucleosome), the one with lower CCC value was removed. Using various CCC cutoffs we calculated established metrics of machine learning(43, 44), namely recall, precision, and F1 (the harmonic mean of precision and recall; see Methods), to compare assigned positions/orientations with the ground truth. Analyzing the nucleosome centers of mass, we obtained a maximum F1 score of 0.85. Adding orientation angles to the comparison reduced the maximum F1 score to 0.76 (Fig S4f). Using the well established template matching program PyTom (41) with default parameters yielded similar results (Fig S4d-g).

**Figure 3.**
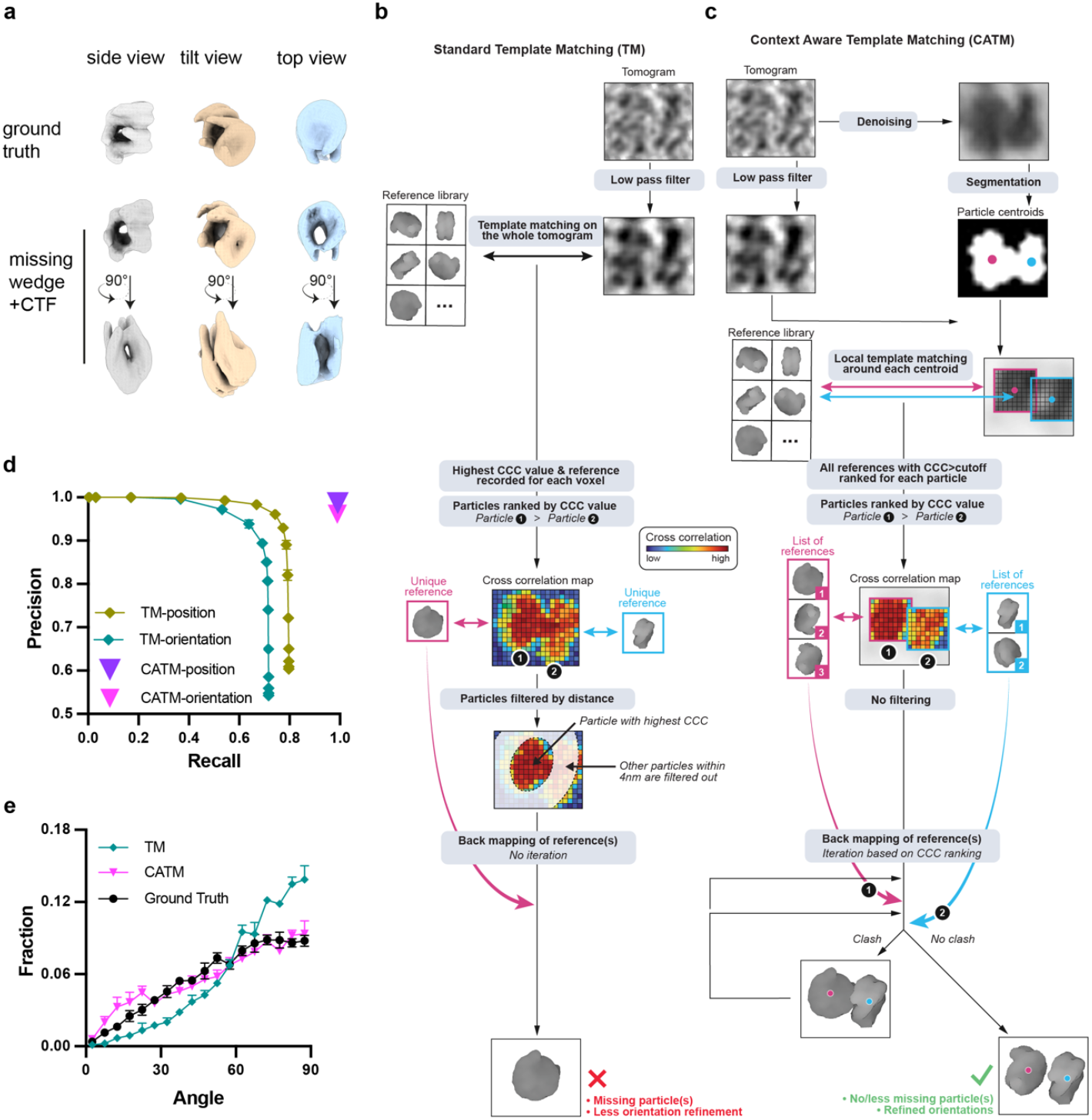
Context-Aware Template Matching for precise nucleosome localization and orientation in simulated data. **(a)** Schematic overview of standard template matching algorithm and context-aware template matching. For standard template matching, each representative voxel only retains the reference image (and corresponding template) with highest CCC to data, and the particles are then filtered by a distance cutoff between their centers of mass. **(b)** Schematic overview of context-aware template matching. Centroids of the models are first identified with deep learning segmentation and localization. The localized particles are then used for local template matching and multiple templates are systematically used for optimization in the assignments. **(c)** Simulation of missing-wedge and CTF modulation effects for a pair of stacked nucleosomes from different perspectives. **(d)** Precision and recall for nucleosome assignments (center of mass position) by CATM and TM. The TM algorithm has different performance characteristics for different cross-correlation cutoffs; the cutoff with highest F1 score was used for later analysis. CATM produces only a single set of assignments. The results are presented as the mean and standard deviation calculated from three tomograms, Note that error bars for CATM are smaller than the symbols. **(e)** Distribution of angle between the normal to the nucleosome plane and the beam direction (Z-axis) as determined by CATM and TM. Nucleosomes are oriented randomly in the ground truth (black circles), producing a sinusoidal distribution. The results are reported as the mean and standard deviation derived from three tomograms.

The high density of particles in condensates often results in overlap of initially assigned positions during template matching, a problem that is exacerbated by artifacts from the missing wedge and CTF (39). To solve this problem, it is essential to simultaneously optimize particle positions and orientations. Moreover, the non-spherical shape of the nucleosomes necessitates that the assignment process incorporates steric information instead of merely relying on a single center-to-center distance cutoff (45). The template matching struggles in this regard because it treats each voxel uniformly, resulting in an unfeasibly large search space when optimizing orientations of multiple particles simultaneously. To address this complexity, we divided the task into two distinct stages: localization and angular assignment (Fig 3c). Initial localization of the particles was achieved using deep learning-based image segmentation (46). This was followed by refinement of particle orientations and positions through local template matching and pose optimization, taking into account the steric properties of the templates (Fig 3c). Local template matching limited by the image segmentation has a vastly limited search space, allowing simultaneous pair-wise optimization of particle position/orientation.

In practice, we used DeepFinder (46) to segment the denoised tomograms (see below) and determine an initial estimate of the center of mass of each particle. We then use these particle positions as a starting point for a process we term Context Aware Template Matching (CATM, Fig 3c, Movie 2). During CATM, subtomograms centered at the particle positions were extracted from the low pass filtered raw tomogram, and the template was rotated at various orientations to calculate the cross-correlation. We retained the orientation with the highest CCC along with those with lower values for each particle (above a threshold of ~0.3, Fig 3c). The mapping process involved iterating through each particle, in descending order of CCC, and positioning them back into a tomogram. If a given particle had a steric clash with previously placed particles, the alternative orientations saved for that particle were examined to resolve the clash. If the clash remained unresolved by these configurations, we paired the nearest neighboring particle with the current particle and tested combinations of orientations for both to settle the conflict. If sterically compatible solutions were found, the orientation pairs that maximized the CCC values were retained and placed in the tomogram. If a resolution was unattainable, only the particle with the higher CCC was saved (Fig S5).

This method significantly improved the accuracy of nucleosome identification and orientation assignments, achieving both high precision and recall (Fig 3d). The F1 score of 0.99 for position assignment and 0.96 for angular assignment (Fig S4e, f) substantially surpasses the TM performance. Furthermore, an important measure of accuracy in the assigned nucleosome orientations is the distribution of angles with respect to the imaging axes. Since nucleosomes should be randomly oriented both in the simulated data and in real data on frozen condensates (which are isotropic), the distribution of nucleosome planes should be random, i.e. sinusoidal in angular coordinates (47) (Fig 3e, black line). Missing wedge distortions make correct assignment in the Z-dimension particularly difficult. As shown in Figures S4g-h, standard template matching under-counts nucleosomes viewed face-on (low angles between the vector normal to the nucleosome plane and the imaging axis vector, along Z) and over-counts side-on views (high angles between these vectors). In contrast, CATM achieved a random distribution of nucleosome orientations that mirrored the ground truth. Thus, image segmentation followed by CATM enables nucleosomes to be accurately positioned and oriented in simulated tomograms.

### Application of CATM to Chromatin Condensates

We next sought to apply CATM to reconstructed tomograms generated from tilt series acquired on cryo-FIB milled lamella containing chromatin condensates. To enhance the visualization and segmentation of nucleosomes, we first used Warp (48) and IsoNet (49) to denoise the tomograms (Fig S6). We manually segmented a small training set volume in DeepFinder, and used the assignments to train the neural network for segmentation. We employed the MeanShift (50) algorithm to retrieve the centroids of the particles and to remove particles too close to the surfaces (< 20 nm) of the FIB-milled lamella or located external to the condensates (Fig S5).

Based on the particle locations, we used CATM to assign nucleosome orientations. Both position and orientation were then further refined using Relion (51). We noticed that since particles are independently processed in Relion, refinement has a propensity to cause nearby particles to become overlaid at a single position. This occurs for ~5-20% of particles depending on the nature of the condensate (Fig S7a-c). To ensure accurate placement of the maximum number of particles, we merge the results from CATM and Relion, excluding overlaid particles that lead to clashes and adding the missing particles from the CATM result, to achieve a refined set of nucleosome assignments (Fig S7a,b).

Visual inspection of the final set of assigned particles mapped back to the tomograms suggested that most nucleosomes were correctly positioned and oriented by the procedures (Fig 4a). Moreover, the distribution of nucleosome orientations was nearly random in the imaging directions, as expected for an isotropic condensate, supporting accuracy of the assignments (Fig 4b). Finally, we extracted subtomograms and performed subtomogram averaging using Relion, which yielded a nucleosome structure at 6.1 Å resolution without discarding any particles (126,126 particles from 14 tomograms acquired from eight lamella, Fig 4c, S8a-b, Supplementary table S1). These qualitative and quantitative measures strongly suggest that our image analysis procedures accurately identify the positions and orientations of most nucleosomes within the dense interior of chromatin condensates. This information enabled us to construct a graph network of nucleosome interactions (Fig. 4d). To assess the heterogeneity of the node valency distribution of the network, we compute the entropy of the distribution (52). The chromatin network exhibits heterogeneity, with a normalized entropy of valency distribution of 0.74±0.02 on a scale of 0 to 1. This analysis reveals that at high resolution (10-100 nm) nucleosomes in the condensates are organized heterogeneously, with small clusters of high density and locally high valence surrounded by regions of low density/valence. Such meso-scale heterogeneity has been observed in computational (53) and experimental (54) studies of condensates produced by intrinsically disordered proteins, suggesting that it may be a general feature of phase separated compartments. Thus, our pipeline from sample preparation to image analysis has enabled us to visualize the internal structure of in vitro reconstituted chromatin condensates, both at the level of individual nucleosome structures and the meso-scale organization of the interaction network.

**Figure 4.**
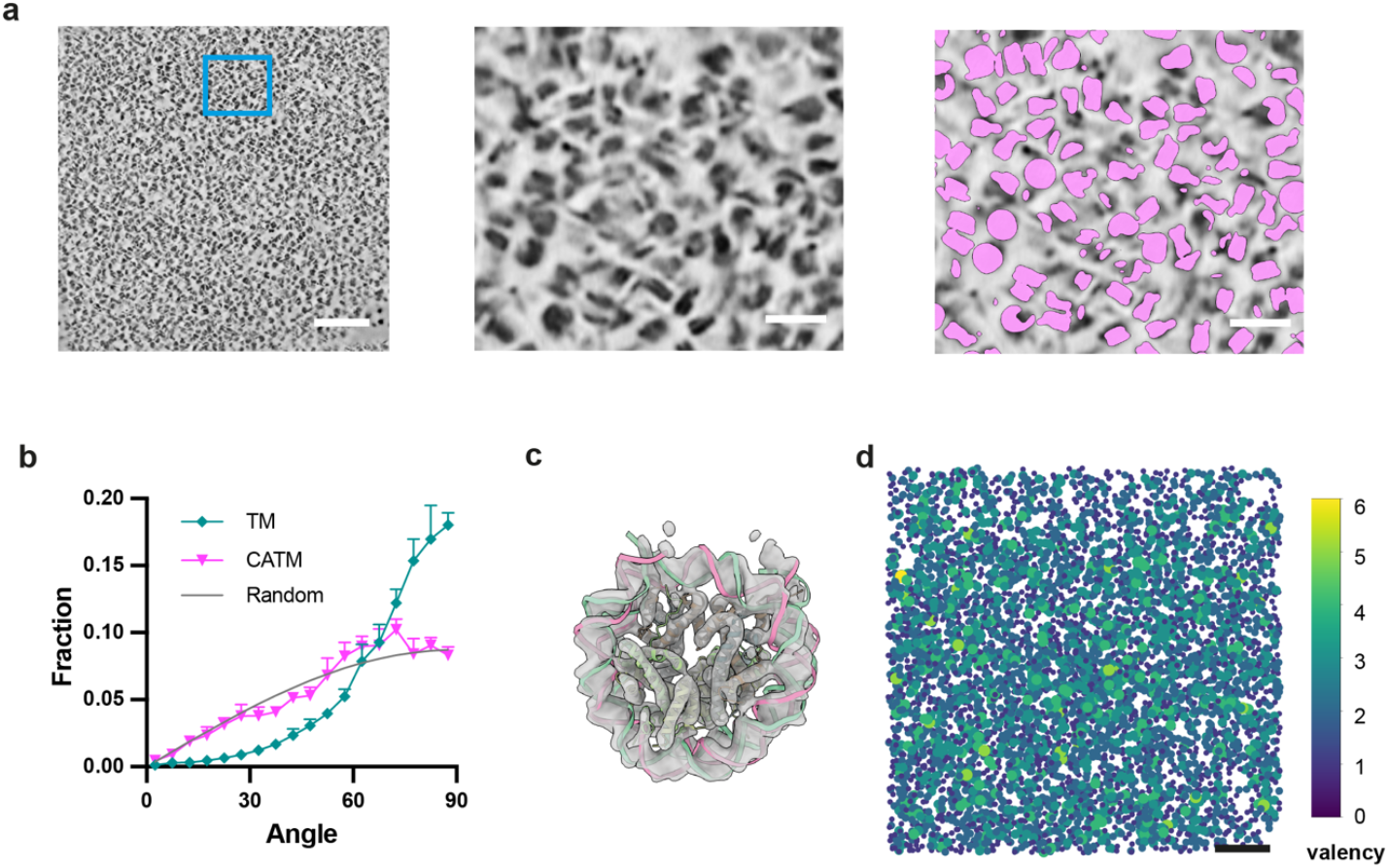
Cryo-ET analysis of reconstituted chromatin condensates. **(a)** Tomographic visualization and CATM analysis of chromatin condensates. Left panel presents a tomographic section through a chromatin condensate, scale bar is 100 nm. The central panel shows an enlarged view of the area delineated by the blue square in the left image, scale bar is 20 nm. The right panel shows CATM nucleosome assignments for the tomographic section in the central panel. **(b)** Distribution of nucleosome orientations with respect to the beam direction (Z-axis) as determined by the specified algorithm. Random orientations (grey) are sinusoidally distributed. **(c)** Subtomogram averaging of 126,125 particles from 14 tomograms, identified by CATM, aligned with a cryo-EM derived core nucleosome structure (PDB ID: 6pwe). **(d)** Graph network of the chromatin condensate, where each node represents a nucleosome from panel (a) and is color-coded based on the valency of interactions it mediates.

### Application of CATM to *In Situ* Native Chromatin

We next sought to apply our methods to native chromatin, which is appreciably more complicated than our synthetic condensates. Complexity arises because in the cell, nucleosomes are composed of different histone variants (55), are covalently modified with diverse epigenetic marks (56), are separated by DNA linkers of variable length (57), and are bound to other macromolecular components (58). Nevertheless, the relatively stereotypical structure of the nucleosome, coupled with previous cryo-ET applications to native chromatin(17, 30–34), suggested our approaches could be effective even in this more complicated system.

We purified HeLa cell nuclei and prepared lamella using HPF and cryo-FIB milling. Consistent with super-resolution imaging of nuclear DNA (59, 60), native chromatin shows numerous 100-300 nm diameter regions of high electron density separated by regions of low density. Within the former, numerous nucleosomes can be observed readily by inspection (Fig 5a). We focused our analyses on these nucleosome-rich regions. These samples yielded high-quality tomographic data, enabling us to precisely map nucleosomes using CATM (Fig 5a). In the absence of specific landmarks that might locally influence the organization of chromatin, we would expect nucleosomes to be randomly oriented, as in our in vitro condensates. Consistent with this idea, nucleosome orientations in native chromatin were predominantly random with respect to the imaging axes (Fig 5b). This random distribution supports both the maintenance of sample integrity during freezing, and the efficacy of our algorithm. Subsequent subtomogram averaging yielded a nucleosome structure with a resolution of 12 Å, based on 35,503 particles observed in three lamellae (Fig 5c, S8c, Supplementary table S1).

**Figure 5.**
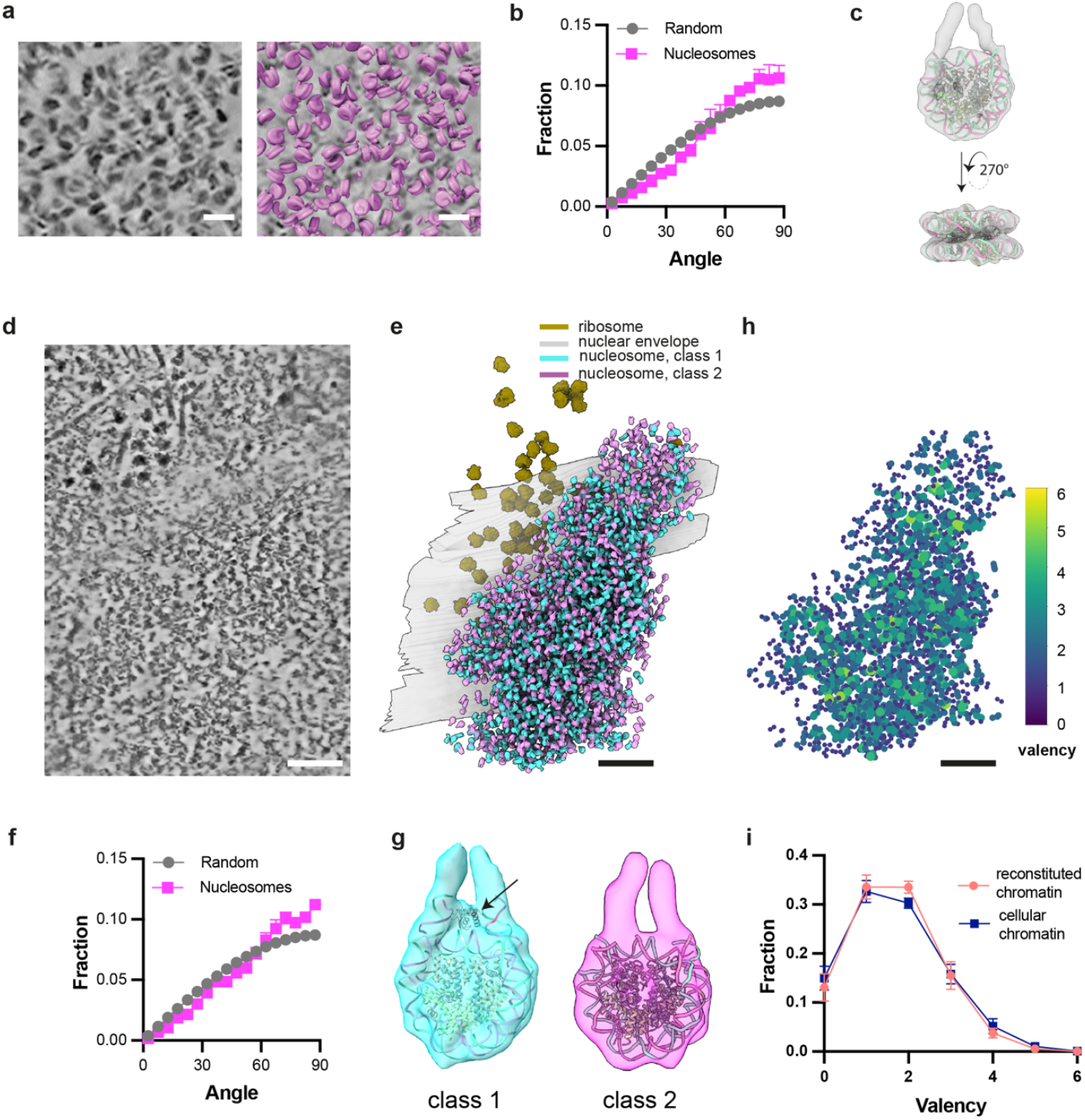
Configuration of native chromatin in isolated HeLa nuclei and intact NIH3T3 Cells. **(a)** A tomographic slice through purified HeLa cell nuclei (left) with superimposed nucleosome models assigned by CATM (right). Scale bar is 20 nm. **(b)** Distribution of angle between the normal to the nucleosome plane and the beam direction (Z-axis) as determined by CATM of the 35,503 nucleosomes identified within 13 tomograms from three purified HeLa cell nuclei. **(c)** Subtomogram average of nucleosome structure extracted from the HeLa cell nuclei with resolution is 12 Å, aligning with nucleosome structure (PDB ID: 6pwe) **(d)** Cross-section from a tomographic volume of an NIH3T3 cell. Scale bar is 100 nm. **(e)** Segmented visualization of the nuclear envelope, ribosome and nucleosomes from the tomogram in (d) but with whole tomogram annotation, illustrating nucleosomes assigned by CATM (magenta and cyan), ribosomes (brown), and the nuclear envelope (gray) embedded within the volume. **(f)** Distribution of angle between the normal to the nucleosome plane and the beam direction (Z-axis) as determined by CATM of 13,265 nucleosomes from four tomograms of NIH3T3 cells. **(g)** Subtomogram average depicting the two classes of nucleosome reconstructed from a total of 6470 and 6804 particles in 4 tomograms from NIH3T3 cells. The resolution is 12 Å for class 1 and 22 Å for class 2, with the maps fitted to nucleosome structures (PDB IDs: 6pwe and 4qlc, respectively). Black arrow points to probable linker histone density at the nucleosome dyad. **(h)** Graph network of the cellular chromatin condensate, where each node represents a nucleosome (not to scale) from panel (a) and is color-coded based on the valency of interactions it mediates. **(i)** The valency distribution of chromatin graph networks in both reconstituted chromatin (illustrated in Fig 4) and cellular chromatin from NIH3T3 cells, presented as the mean and standard deviation derived from four tomograms.

We further extended our approaches to plunge-frozen NIH3T3 cells, where we collected tilt-series on cryo-FIB milled lamella containing nuclei. Within the peripheral regions of these nuclei, we identified various cellular components, including ribosomes, the nuclear envelope, and nucleosomes. Post-denoising, regions containing nucleosomes at high density were clearly distinguishable (Fig 5d). Within these regions, individual nucleosomes could be assigned using CATM (Fig 5e). The near-random distribution of nucleosome orientations relative to the imaging axis confirmed the accuracy of the algorithm and indicates that in vivo the higher order organization of chromatin may not be strongly influenced by neighboring cellular structures (Fig 5f). Subtomogram averaging yielded two different classes of nucleosome structures, with ~12 Å and 22 Å resolution, respectively (Fig 5g, S8d, Supplementary table 1). Notably, in the former, the reconstructed structure included density consistent with linker histone, which binds to the nucleosome dyad axis with high stoichiometry in mammalian cells (61) (Fig 5h).

In contrast to the reconstituted chromatin condensates, where the entry and exit DNA of nucleosomes was not visible in reconstructions (Fig 4c), in both of the cellular samples this DNA is readily observed (Fig 5c, g), suggesting less conformational variability. The more limited conformational space in cells could arise from several factors including DNA linker lengths, nucleosome binding molecules, different histone variants, or histone post-translational modifications and/or their reader proteins. Further studies correlating nucleosome organization and structure with these factors will be necessary to understand these observations.

Finally, constructing graph networks of the cellular chromatin revealed that like the reconstituted condensates, nucleosomes are organized heterogeneously with normalized entropy of valency distribution of 0.766±0.009 (versus 0.74±0.02 for chromatin condensates, see above and Fig 4d). Small foci of high local density and valency are interspersed within larger regions where both parameters are lower (Fig 5h). The similarity of the networks found in native chromatin and synthetic condensates suggests that cellular chromatin fibers, despite their vastly longer lengths, have sufficient plasticity to pack analogously to much smaller fragments. The similar packing in cells and in reconstitutions likely arises from the relatively stereotypical structure of nucleosomes and the rigidity of linker DNA. These characteristics of chromatin may be analogous to the fact that within folded proteins, amino acids adopt packing and geometries that are similar to those found in amino acid crystals (62).

## Discussion

There is growing recognition that biomolecular condensates have highly complex physical properties (63). Most behave as heterogeneous network fluids, with internal substructures on different length scales (64), as well as differences in molecular organization between their cores and surfaces (65, 66). Moreover, their internal solution environments can differ substantially from the surrounding media, for example in pH, salt concentration (67, 68), and hydrophobicity (69). To fully understand how these properties affect chemistry within condensates, it is necessary to understand the structures of individual components as well as their higher order organization.

Our results demonstrate the effectiveness of high-pressure freezing (HPF) and focused ion beam (FIB) milling (the waffle method (37)), coupled with cryo-ET and image analysis, to achieve this goal. The pipeline we have developed here overcomes two substantial obstacles in studies of condensate structure. First, due to the liquid-like properties of condensates, inappropriate preparation of biochemically reconstituted samples can lead to artifactual distortions of both their overall architecture (Fig. 1b) and their internal packing arrangements (Fig 1e, S1e, S2b). By contrast, HPF is highly structure-preserving. Second, the high density of condensates, both biochemically reconstituted and cellular, can lead to inaccurate molecular assignments, introducing artifacts such as apparent preferred orientation (Fig 3, 4b, S4d-h), which impair interpretation of higher order organization and can degrade the quality of subtomogram analysis. Our CATM algorithm utilizes both features of the images and prior knowledge of molecular structure to achieve accurate assignments. We note that while artifacts of sample preparation and image analysis are readily identifiable in chromatin condensates due to the nature of nucleosomes (large, stereotypical structure; high electron density), they may not be as obvious in other types of condensates. Nevertheless, they are expected to occur in cryo-ET studies of many liquid-like systems of high density, emphasizing the importance of the workflow we have developed here. These methods now open the door to understanding how the internal structure of biomolecular condensates influences diverse processes.

Further technical improvements should advance the capabilities of the pipeline. Recently matured plasma-FIB milling (70) will enhance the efficiency of sample preparation, as it can process thick ice layers more rapidly and potentially yield more stable lamellae. Additionally, for condensates typically less than 10 µm in diameter, employing specially designed grids with thinner bars could reduce stress on the samples during milling, leading to quicker processing and more stable lamellae. Moreover, advanced laser phase plate technology (71) holds great promise for visualizing and analyzing smaller, more heterogeneous samples in both in vitro and in vivo settings. Our CATM algorithm may be enhanced by incorporating GPU acceleration, which could improve angular sampling during template matching, thereby refining the precision of angular assignments. Moreover, expanding from two-body to multi-body optimization, possibly incorporating Monte Carlo sampling, could provide even more accurate assignments, advancing our understanding of complex interactions within condensates.

Human cells compact 2 meters of DNA into a 10 µm nucleus by wrapping DNA around histones to form nucleosomes, which assemble into higher-order structures and interact with proteins to regulate gene transcription and cell fate. A key question is how the structures, interactions, and 3D organization of chromatin regulate cellular processes across multiple scales. Studies have revealed that cellular chromatin forms small clutches (~10–50 nucleosomes) (72, 73) and larger Topologically Associated Domains (TADs, ~1000-5000 nucleosomes), but the precise positions and orientations of nucleosomes within these assemblies remain elusive due to technical challenges (73, 74). While selective DNA labeling improves visualization of chromatin at finer scales (75), it often sacrifices structural details. Cryo-EM offers great potential to observe chromatin in its native state, particularly when samples are generated with high-pressure freezing (HPF) combined with cryo-FIB milling (34, 37, 76). Reliable preparation of native samples, coupled with advanced quantitative analysis, could enable precise probing of chromatin network interactions and their functional implications. Furthermore, advanced algorithms capable of automatically tracing nucleosome chains in chromatin could provide insight not only into the conformations of small clutches but also the connectivity within TAD-like domains.

Most of the cryo-ET analyses described here focused on biochemical reconstitutions, where knowledge of component parts greatly simplifies template matching. Nevertheless, we have shown that our pipeline can be extended to certain cellular structures where selected components are highly enriched and have stereotypical structures, as in chromatin studied in nuclei here. Other cases may include condensates containing recognizable filaments, as in centrosomes (77) or immune signaling condensates (10), or large machines, as in transcriptional (78) or splicing foci (79). Advances in correlative cryo-super-resolution light microscopy and cryo-ET may enable molecular-scale structural analyses of other condensates as well (80). Ultimately, the combination of biochemical reconstitutions of increasing complexity (7, 10, 77) plus in situ analysis of cellular condensates is likely to provide the greatest depth of mechanistic understanding.

In conclusion, by ensuring the integrity of reconstituted condensates and accurate molecular assignments, our study establishes a workflow for structural analysis of both biochemically reconstituted and native chromatin condensates, which should be applicable to certain other condensates as well. Using it, we have been able to determine structures of nucleosomes in both reconstituted chromatin condensates and native chromatin. Additionally, we found that in both cases that higher order nucleosome packing is heterogeneous. Future studies will address key issues in the condensate and chromatin fields, including mechanisms of phase separation, the activities of resident molecules, and the influence of environment on condensate chemistry

## Acknowledgements

We thank the following imaging facilities at UT southwestern medical center : cryoEM microscopy facility (CEMF, CPRIT Core Facility Support Award RP220582), Structure Biology Lab (SBL, CPRIT Core Facility Support Award RP220582), Electron microscopy core facility (EMCF), BioHPC. We thank the training course at the National Center For In-Situ Tomographic Ultramicroscopy (NCITU) supported by the NIH Common Fund Transformative High Resolution Cryo-Electron Microscopy program (U24 GM129539). Research was supported by the Howard Hughes Medical Institute (M.K.R and E.V.), a Paul G. Allen Frontiers Distinguished Investigator Award (to M.K.R.), grants from the Welch Foundation (I-1544 to M.K.R.), the National Institutes of Health (R35GM141736 to M.K.R.), and EMBO postdoctoral fellowship (ALTF 871-2020 to J.H.) We thank Daniela Nicastro for mentoring and advice at the early stage of this project.

## Code availability

The TM and CATM software, and all analysis scripts are available on Gitlab: https://git.biohpc.swmed.edu/rosen-lab/catm

## Data availability

All data in the manuscript will be uploaded to EMDB before publication, and will be provided to referees upon request.

## Author contributions

Conceived of the study, H.Z., M.K.R. Generated biochemical samples, H.Z., L.K.D.

Generated cryoEM samples, M.S., H.Z., N.J., J.H.

Collected cryoEM data, H.Z., X.Z, S.Y., R.Y., Z.Y., J.H

Developed cryoET analysis methods, H.Z.

Developed nucleosome orientation analysis methods, H.Z., J.H.

Supervised project, E.V., Z.Y., M.K.R.

Drafted manuscript, H.Z.

Generated illustrations and Movies, M.R.

Revised manuscript, all authors

## Materials and Methods

### Protein purification and nucleosome array assembly

The nucleosome arrays were constructed as previously described (23). Briefly, human histones H2A, H2B, H3, and H4 were recombinantly expressed in E. coli, purified chromatographically and assembled into octamers. Octamers were assembled using salt dialysis onto DNA consisting of 12 repeats of the Widom 601 nucleosome positioning sequence. Labeling was performed by incorporating 1% of AF594-labeled H2BT116C octamers during assembly. A 25% excess of H2A/H2B dimer was added to facilitate complete octamer loading. DNA fragments of ~300 bp were used as carrier DNA to absorb the excess histones to avoid over assembly. The assembled nucleosome arrays were purified using sucrose gradient centrifugation. Sucrose was removed by dialysis and the chromatin was then concentrated by ultra-filtration to 6-8 µM in 20 mM Tris-HCl, pH 7.5, 1 mM EGTA, 1 mM DTT. Array quality was assessed by micrococcal nuclease digestion into mononucleosomes, followed native polyacrylamide gel electrophoresis (PAGE). Only samples with >90% properly assembled octamer (rather than under-assembled hexamer, lacking one copy each of H2A and H2B) were analyzed further.

### Purification of HeLa Cell Nuclei

HeLa cell pellets (5×109 cells, Ipracell) were resuspended in 5 volumes of lysis buffer (20 mM HEPES, pH 7.9, 5 mM Mg(OAc)2, 0.15 uM Spermine, 10 mM Sodium Butyrate, 10 mM Nicotinamide, 10 uM Pepstatin, 3 mM AEBSF, 10 uM E-64, 2 uM Antipain, 500 uM leupeptin, 2ug/ml aprotinin, 4 uM TSA, 1 mM DTT), and swelled on ice for 10 minutes. Cells were lysed with 0.05% NP-40 and after gentle inversion were kept on ice for 10 minutes. The lysate was centrifuged at 1000 x g for 10 minutes at 4°C to pellet the nuclei. The supernatant was carefully removed, and the nuclear pellet was washed twice with lysis buffer containing 0.34 M sucrose by resuspension and subsequent centrifugation at 1000 x g for 10 minutes to remove cytoplasmic contaminants. The integrity and purity of the isolated nuclei were verified by light microscopy. Purified nuclei were flash frozen in liquid nitrogen and stored at −80 °C. For HPF experiments, frozen nuclei were resuspended in freezing buffer (20 mM HEPES, pH 7.9, 100 mM KOAc, 1 mM Mg(OAc)2, 0.15 uM Spermine, 10 mM Sodium Butyrate, 10 mM Nicotinamide, 10 uM Pepstatin, 3 mM AEBSF, 10 uM E-64, 2 uM Antipain, 500 uM leupeptin, 2ug/ml aprotinin, 4 uM TSA, 1 mM DTT).

### Grid preparation

#### Blotting method

For sample preparation using the Vitrobot Mark IV (Thermo Fisher), Lacey carbon grids (200 mesh, EMS) were glow discharged at 30 mA for 30 seconds prior to use. Nucleosome arrays were first equilibrated in a dilute buffer (20 mM Tris-OAc, pH 7.5, 0.1 mM EGTA) at a concentration of 1 µM. They were then mixed 1:1 with buffer containing 40 mM Tris-OAc, 150 mM KOAc, 2 mM Mg(OAc)_2_, and 0.2 mM EGTA to induce phase separation. The droplets formed instantly and solutions were transferred to grids within 10 minutes. Blotting was conducted at 4 °C and 100% humidity. 3 µL of chromatin solution was applied to the grids, blotted with force 0 for 3.5 seconds, and plunge frozen in liquid ethane, yielding samples with 60 nm to 120 nm thickness.

For single-side blotting, we used an in-house manual plunge freezer. The sample and grids were prepared as described above, with chromatin droplets added to the frontside of the grid. The grid was blotted with filter paper from the backside for 3.5 seconds and then plunge-frozen in liquid ethane.

#### Self-wicking method

Utilizing the Chameleon system (SPT Labtech) for the self-wicking method, nucleosome arrays were pre-equilibrated in a dilute buffer (20 mM Tris-OAc, pH 7.5, 0.1 mM EGTA) at 0.5 µM concentration, mixed 1:1 with buffer containing 40 mM Tris-OAc, 200 mM KOAc, 2 mM Mg(OAc)_2_, 0.2 mM EGTA to induce phase separation. Within 10 minutes, 5 µl of the phase-separated chromatin solution was applied to self-wicking grids, which had been glow discharged at 12 mA for 20 seconds using the internal glow discharger. A strip of liquid was sprayed onto the grid, allowed to wick for 120 milliseconds before plunge freezing, yielding samples with 80 nm to 160 nm thickness.

#### Waffle method

Prior to sample preparation, planchets for the high-pressure freezer were pre-treated with 0.05% lecithin and Quantifoil carbon 200 mesh EM grids (Quantifoil) were glow discharged at 15 mA for 30 seconds. Lecithin treatment proved less contamination than other solvents (e.g. hexadecane) in affording flat vitreous ice surfaces of the slab following freezing. For chromatin condensates, the nucleosome arrays were pre-equilibrated in a dilute buffer at 6 µM concentration, then mixed with buffer containing 40 mM Tris-OAc, 200 mM KOAc, 2 mM Mg(OAc)_2_, 10% glycerol, 0.2 mM EGTA to induce phase separation. After placing the first planchet on the HPF tip holder, the EM grid was placed on the planchet with the back side facing up, 4 µL of the phase separated chromatin sample was applied (within 10 minutes of inducing phase separation) to the EM grid, and allowed to sit for 30 seconds before assembly of the HPF tip holder. Settling time was an important optimization parameter to ensure a sufficient density of condensates near the vitreous ice surface to afford efficient targeting during subsequent FIB milling. The sample-loaded grids were inserted into a Wohlwend HPF Compact 03 machine, where rapid freezing was conducted at 2050 bar using liquid nitrogen at −196°C. Following freezing, the grids were disassembled from the HPF tip and stored in liquid nitrogen until required for cryo-fluorescent imaging. For the purified nuclei, the sample was frozen using the same procedure as the reconstituted chromatin.

#### Plunge freezing of NIH3T3 cells

NIH-3T3 cells (ATCC CRL-1658) were maintained at 37 °C, 5% CO_2_ in DMEM (Gibco 11995073). Approximately 1.25×10^5^ trypsinized cells were seeded onto glow-discharged EM grids (carbon-coated Quantifoil R1/4 Au200-C) and allowed to adhere for 2 hours at 37 °C, 5% CO_2_. Cells were treated with 100 nM jasplakinolide for two hours. Grids were manually blotted with Whatman filter paper #1and plunged in liquid ethane/propane (50:50, AirGas) using a custom-built plunging and vitrification device (Max Planck Institute for Biochemistry, Munich).

### Correlative Cryo-Light and Electron Microscopy (CLEM) for Condensate Samples

Chromatin droplet cryo-grids were examined using a Leica cryo-thunder CLEM system equipped with a 50x/dry objective (NA=0.9). For fluorescence detection, TXR filter cubes (emission wavelength: 592-668 nm) were utilized. The fluorescence intensity manager was set to 100% with an exposure duration of 600 ms. To comprehensively image the entire grid, an 8×8 tile array encompassing a z-stack of 10-20 μm with a 0.5 μm step increment was assembled using Leica LAS X software. These z-stack montages were then processed into 2D CLEM images via maximum intensity projection, facilitating subsequent correlation with cryo-FIB milling processes.

### Cryo-FIB-milling

#### HPF samples (Reconstituted chromatin and purified nuclei)

Lamellae were milled using the waffle method (37, 81)followed by manual polishing. Grids were loaded onto a pre-tilted shuttle (45° or 35°), then transferred into an Aquilos 2 cryoFIB-SEM (Thermo Fisher Scientific). The stage temperature was maintained below –180 °C. A grid overview SEM image was taken at 2 kV, 13 pA, then an image of the grid center was acquired with a higher voltage (5-20 kV, 13 pA) to visualize the center landmark of the grid. A CLEM image was imported to Maps (v.3.16-3.25) and then aligned to the SEM image by using the center landmark shape of the grid. Squares with abundant fluorescent droplets were picked and subregions with dense droplets were targeted during the milling. The grids were sputter coated with platinum (30 mA, 10 Pa, 15s) then an organometallic platinum layer was deposited by a gas injection system for 2.5-3 minutes.

Precuts were milled following the waffle method protocol (37, 81) with a beam current of 7-15 nA at 30 kV. Next, a secondary organometallic platinum GIS layer was deposited for 2.5-3 minutes to ensure sufficient GIS layer left through the following procedure. An updated SEM image was taken to highlight the locations of the precuts in Maps software. AutoTEM (v.2.0-2.3, ThermoFisher Scientific) was used to define eucentric position and milling position for each lamella site, then the underside of the potential lamella was manually milled using a beam current of 3-5 nA at three incrementally decreasing milling angles; 40°, 30°, 20° for a 35° pre-tilted shuttle, or at two angles; ~24°, 20° for a 45° pre-tilted shuttle. SEM images were taken to inspect and monitor the lamella to avoid double layers. At the 20° milling angle, a notch pattern (37, 81) was milled using a beam current of 0.3 nA for 2.5 minutes. AutoTEM was then used to automate the milling at each lamella site, aiming for a final thickness of 180-300 nm at 20° milling angle. The lamella width was set to 12 µm to balance the stability and size of the lamella. The FIB-milling parameters described in the waffle method (37, 81) were used with adjustments of depth correction.

After AutoTEM milling, the lamellae were manually polished to ~100-150 nm, utilizing stage over-tilts of +0.5° (20.5°) and –0.2° (19.8°) from the original milling angle (20°). A rectangle pattern or cleaning cross section (CCS) was used to polish the lamella with a beam current of 10-50 pA. For CCS pattern, Z size was set to 1 µm so the lamella is not damaged during milling. Lastly, the stage was returned to the original milling angle of 20° for a final polish. We note that lamella thickness < 150 nm was essential to achieving high quality cryoET data that were sufficient for accurate template matching subsequently.

#### Plunge frozen samples (intact NIH3T3 cells)

Grids were milled using an Aquilos 2 cryoFIB-SEM (Thermo Fisher Scientific) using a combination of automated milling (AutoTEM) and manual polishing with 10 pA. Briefly, milling progressed from rough to fine milling using currents at 500, 300, 100, 50, 30 and 10 pA aiming for a nominal lamella thickness of 150 nm. Lamella were polished at 10 pA with 0.5° overtilt for approximately two minutes to remove thicker material from the back of the lamella.

### Cryo-ET Data Acquisition

#### Blotting and Self-wicking Samples

Data acquisition for samples prepared by blotting and self-wicking was carried out using a Titan Krios G1 (Thermo Fisher Scientific) equipped with a K3 camera (Gatan, Inc) operating at 300 kV. Tilt series were captured ranging from −60° to +60°, with a 3-degree increment per tilt. Images were recorded at a pixel size of 0.206 nm at the specimen level. A Volta phase plate was employed, and the defocus was set to −0.5 µm. To minimize radiation damage, the total electron dose was limited to 150 electrons/Å^2^.

#### Condensate lamellae samples and purified nuclei

For condensate lamellae, data acquisition was conducted using a Titan Krios G3 (Thermo Fisher Scientific) featuring a cold-field emission gun, a Selectris X imaging filter, and a Falcon 4i camera. Tilt series ranged from −48° to +60° with a 2-degree increment per tilt. Each image was captured at a physical pixel size of 0.1516 nm at the specimen level. The defocus ranged between −3 µm and −4.5 µm, with a total electron dose capped at 178 electrons/Å^2^.

#### Cellular samples

Milled grids were imaged on a Titan Krios G3 microscope (Thermo Fisher Scientific) operated at 300 keV equipped with a K3 detector and 1067HD BioContinuum energy filter (Gatan) with 15 eV slit-width. Dose-fractionated images were acquired using SerialEM(82). Dose-symmetric tilt series were acquired in low dose mode using the parallel cryoelectron tomography (PACEtomo) scheme (83) with 3° increments and +/-54° tilt range from a 6-12° lamella pre-tilt. A total dose of approximately 140 e/Å2 at 1.34 Å/pixel and 4 µm nominal defocus was used (Supplementary Data Table 1).

### Tilt-series Processing and Alignment

The acquired movie frames underwent gain correction, motion correction and Contrast Transfer Function (CTF) estimation using Warp (48). Subsequent tilt series creation and alignment were performed in AreTomo(84). These alignments facilitated further processing in Warp to reconstruct tomograms at 8 Å/pixel. Additionally, two half-sets tomograms were constructed to enable denoising.

### Tomogram denoising and segmentation

Initially, the two half-sets of tomograms along with their respective CTF models were used to train a denoising model in Warp, undergoing 40,000 iterations. This process significantly enhanced the contrast of the tomograms. Subsequently, the denoised tomograms were processed through IsoNet for an additional round of missing-wedge restoration and further denoising. The models were specifically retrained for each dataset to ensure optimal performance.

For segmentation, a small section of the tomogram measuring 256×256×304 voxels was manually annotated to identify nucleosome-containing voxels, which served as the training set for DeepFinder. Following training of the neural network, the full tomograms were segmented to identify voxels representing nucleosomes.

The centroid of each particle was then determined using MeanShift, which clustered the segmented maps within a window size of 4.5 pixels. To avoid artifacts from the gallium ions used in the milling process (85), centroids located less than 20 nm from the FIB-milled surface were removed from the dataset. Due to imperfections in the segmentation, regions with a noisy background outside of the condensate sometimes were also picked. We used an intensity threshold to remove such false positive particles.

For native chromatin we used WARP/IsoNet to denoise the tomograms, and DeepFinder/MeanShift to segment the images for nucleosomes as above. Then we used IMOD to manually delineate regions with a high density of nucleosomes, and only these were used for downstream analysis. To annotate nuclear envelopes, segmentation was performed in IMOD and a custom script was used to generate a binary mask. Ribosome positions and orientations were sequentially determined using Napari(86) and CATM, respectively.

### Context-aware template matching (CATM)

To enhance molecular assignment within crowded condensates, we developed a context-aware template matching (CATM) procedure. Following deep learning-based particle identification, CATM performs local template matching and clash resolution to accurately determine molecular positions and orientations. For clarity, in the description below we use the following terms: “particle” for a generic object placed at a position in the map, “model” for the low resolution filtered structure or experimentally determined structure, “template” for a model rotated to a particular orientation (used in clash resolution), and “reference” for the image of the template distorted according to missing wedge and CTF effects (used in CCC calculation).

#### Template generation

In the CATM pipeline, tomograms reconstructed in Warp through back projection were low-pass filtered with Gaussian filter at 25 Å using EMAN2(87), and a corresponding CTF model for each tomogram was generated. An initial nucleosome model was generated by low-pass filtering a nucleosome structure (pdb 6pwe) to 25 Å resolution with 8 Å/pixel to match the experimental data. The model was subsequently replaced with an experimental structure determined by subtomogram averaging to achieve better fitting for all the results reported here. Missing wedge and CTF artifacts severely distort the nucleosome structure in tomograms (Fig. 3c, S3). It was essential to properly account for these distortions during template matching, so that the references better represented the real data. We took two steps in this regard. First, the nucleosome model was applied to the 3D CTF model generated from Warp to assess the distortion and choose the contour level of the distorted template to minimize the artifacts. In addition, for a given nucleosome model we created a library of templates representing different orientations of the nucleosome, uniformly sampled in 3D space. Each template was then appropriately corrected for the missing wedge effect and CTF modulation to produce a library of references for use in CCC calculations during template matching as described below.

#### Clash resolution

Following template matching, the list of particles was ranked by their maximum CCCs, and then mapped back to the tomogram sequentially starting with the template whose reference afforded the highest CCC for the particle. In cases where a clash occurred during new particle placement, the other templates for the new particle (corresponding to references with CCC values less than the maximum) were then used to resolve the clash. If the clash could be resolved by another template, the two templates were then saved in an assigned list and mapped to an assigned map. Otherwise, a mechanism was employed to resolve the clash that began by retrieving the nearest neighbors of the pair among the currently mapped particles. The densities of the nearest neighbors were then erased from the assigned map, and a subregion of the assigned map (80 voxels) was subtracted from the assigned map to accelerate template optimization. Within this dense environment, all possible combinations of templates corresponding to both particles were sampled to find a clash-free solution whose references afforded the highest combined CCC value. If a clash could not be resolved—typically due to false positives in segmentation—the particle with the highest CCC value was retained. The process of particle placement and clash resolution continued until all particles identified by segmentation had been accounted for. Note that in addition to placing particles in order of CCC values, we also examined random ordering or spatially defined ordering. Both of these alternatives yielded less random nucleosome orientation distributions than the CCC-based procedure, indicating poorer performance.

The process concluded with the generation of various electron microscopy (EM) format coordinate files, alongside an assigned particles tomogram file, documenting the final particle placements and orientations.

### Benchmarking of Template Matching Algorithms

To evaluate the efficacy of various template matching algorithms, we employed cryotomosim (40) to simulate tomograms, using a nucleosome structure (PDB 6pwe) low-pass filtered to 25 Å. This structure was randomly rotated and embedded in a 3D volume alongside DNA fragments of 25 base pairs, equivalent in number to the nucleosomes, to mimic the environment of experimental tomograms. Vitrified ice simulated effects were added, and tilt series were created under specific conditions(40): 300 keV voltage, spherical aberration of 2.7, sigma of 0.9, defocus of −4 µm, total dose of 150 electrons/Å2, symmetric tilt pattern, pixel size of 8 Å, and a tilt range from −60° to 48° with 2° increments. Subsequent tomograms were reconstructed for analysis.

A soft mask with two voxels of extension beyond the density of the nucleosome template, plus two voxels of soft edge decay was created in Relion and systematically used across all tested software.

#### Standard Template Matching (TM) Algorithm

Reference images corresponding to templates at various angles, and distorted with missing wedge effects, were matched against the tomogram using the same cross-correlation function as employed in CATM. The highest CCC value and its corresponding orientation for each voxel in the tomogram were recorded. Particles were ranked based on CCCs and filtered to remove those within 5 voxels (4 nm) of each other. Remaining templates were mapped back to the tomogram, removing any that clashed (defined by two templates occupying the same voxel in the tomogram). Various CCC cut-offs were used to define particle sets, which were then used to calculate metrics for the benchmark, including precision, recall and F1 score (defined below) by comparison to the ground truth, and angular distribution.

#### Context-Aware Template Matching (CATM)

For CATM testing, neural network weights obtained from real data segmentation with DeepFinder were used to segment the simulated data without additional training. The segmented map provided centroids for particle identification, which were then used in CATM as described above.

#### Template Matching in Pytom

Version 0.971 of Pytom was applied to the same template and mask as TM and CATM. The sampling angular interval was set to 12.85°, and wedge correction was applied to the template. Other parameters were used at their default values. Templates were filtered by a 5-pixel distance cut-off in Pytom, and only non-clashing templates were saved for analysis.

#### Performance metric calculation

Performance was measured for each approach using mechanisms similar to the Shrec 2019 cryo-ET Classification benchmark (44). For positional accuracy, a particle was considered a True Positive (TP) if its predicted centroid was within 4 nm of the ground truth. For orientation accuracy, a vector perpendicular to the nucleosome plane was defined in the reference frame, and orientation was considered a TP if the angle between the predicted and ground truth vectors was < 30°. True Positives (TP) were defined as particles correctly assigned; False Positives (FP) as particles assigned by the program but not present in the ground truth; and False Negatives (FN) as ground truth particles not assigned by the program. Precision was calculated as TP/(TP+FP), recall as TP/(TP+FN) and the F1 score as the harmonic mean of precision and recall.

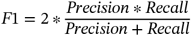

### Subtomogram averaging using Warp-Relion-M

Following CATM, the coordinates and orientation data of the particles were used to reconstruct subtomograms at an 8 Å/pixel resolution in Warp. These subtomograms were subsequently averaged and assembled into an initial model, which was then low pass filtered to 40 Å and used as a reference for further refinement in Relion. Particle refinement involved enhancing precision iteratively while down sampling to 6 Å/pixel, 4 Å/pixel, and finally 2 Å/pixel. This progressive refinement ensured increased resolution and accuracy of the particle models. After completing the final refinement step, the particles were imported into M software(15) for further pose and contrast transfer function (CTF) refinement. This multi-step process leverages the strengths of Warp for initial reconstruction and Relion for detailed refinement, culminating in M for final adjustments, thus optimizing the accuracy and resolution of the resulting structural models.

For the reconstituted chromatin samples and isolated HeLa cell nuclei, averaging was performed for all assigned nucleosomes by CATM; none were discarded. For the NIH3T3 cells sample, the particles were then separated into two different classes, one giving 12 Å resolution for 6470 particles, and a second giving 22 Å resolution for 6804. Resolution estimation was performed with Fourier shell correlation in Relion.

### Analysis of Vitrobot and Chameleon AWI

The air-water interface (AWI) was manually defined for each tomogram with control points using IMOD (88). These were used to create a 2D triangulation mesh using MATLAB and Dynamo scripts (42). For each nucleosome particle, a distance and relative orientation to the AWI was calculated. The relative orientation was taken as the absolute value of the dot product between the surface normal of the closest AWI mesh face and a vector normal to the nucleosome face. To assess preferred orientation at the AWI, the orientation of nucleosomes within 20 nm of the AWI were assessed relative to those further than 20 nm. Oriented nucleosomes were visualized at their native positions within the tomogram using ChimeraX (89) and the ArtiaX plug-in (90).

### Analysis of the nucleosome orientation distribution

In our coordinate system the origin is set to the centroid of the reference nucleosome, which was approximated as having 2-fold symmetry about the dyad axis (i.e. composed of palindromic DNA); the Z-axis is perpendicular to the nucleosome plane, the X-axis extends from the centroid to the nucleosome dyad, and the Y-axis is orthogonal to the X-Z plane. For each nucleosome assigned by template matching we determined the angle between its Z-axis and the beam direction or the two beam-normal directions. Since the angle between a random vector and a plane follows a sinusoidal distribution (47) a random arrangement of nucleosomes will exhibit a similar sinusoidal distribution of these angles.

### Reconstruction of the condensate graph network and calculation of the entropy of valency distribution

To construct the nucleosome networks, particle coordinates were imported using an in-house script, and the geometric graph network was reconstructed using networkX (91). Each particle was treated as a node, and edges were assigned between nodes when the distance between nucleosomes was less than 12 nm. The entropy of valency (H in bits) distribution measures the diversity of the degree distribution (52). It is defined as:

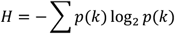

Where p(k) is the probability of a node having degree k. The maximum possible entropy (Hmax) depends on the number of degree values K in the network and for a network with K possible degrees:

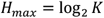

The normalized entropy (H_normalized_) is calculated by normzlize with Hmax for comparision.

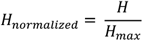

Four tomograms were used to calculate the average and standard deviation of the H_*normalized*_.

## References

1. S. F. Banani, H. O. Lee, A. A. Hyman, M. K. Rosen, Biomolecular condensates: organizers of cellular biochemistry. Nature Reviews Molecular Cell Biology 18, 285–298 (2017).

2. A. S. Lyon, W. B. Peeples, M. K. Rosen, A framework for understanding the functions of biomolecular condensates across scales. Nature Reviews Molecular Cell Biology 22, 215–235 (2021).

3. Y. Shin, C. P. Brangwynne, Liquid phase condensation in cell physiology and disease. Science 357, eaaf4382 (2017).

4. C. Mathieu, R. V. Pappu, J. P. Taylor, Beyond aggregation: Pathological phase transitions in neurodegenerative disease. Science 370, 56–60 (2020).

5. S. Mehta, J. Zhang, Liquid-liquid phase separation drives cellular function and dysfunction in cancer. Nat Rev Cancer 22, 239–252 (2022).

6. D. M. Mitrea, M. Mittasch, B. F. Gomes, I. A. Klein, M. A. Murcko, Modulating biomolecular condensates: a novel approach to drug discovery. Nat Rev Drug Discov 21, 841–862 (2022).

7. S. L. Currie et al., Quantitative reconstitution of yeast RNA processing bodies. Proceedings of the National Academy of Sciences 120, e2214064120 (2023).

8. S. L. Currie, M. K. Rosen, Using quantitative reconstitution to investigate multicomponent condensates. RNA 28, 27–35 (2022).

9. L. B. Case, M. De Pasquale, L. Henry, M. K. Rosen, Synergistic phase separation of two pathways promotes integrin clustering and nascent adhesion formation. Elife 11 (2022).

10. X. Su et al., Phase separation of signaling molecules promotes T cell receptor signal transduction. Science 352, 595–599 (2016).

11. L. N. Young, E. Villa, Bringing Structure to Cell Biology with Cryo-Electron Tomography. Annual Review of Biophysics 52, 573–595 (2023).

12. L. E. Coronas et al., Stability and deformation of biomolecular condensates under the action of shear flow. The Journal of Chemical Physics 160, 215101 (2024).

13. B. A. Gibson et al., In diverse conditions, intrinsic chromatin condensates have liquid-like material properties. Proceedings of the National Academy of Sciences 120, e2218085120 (2023).

14. D. Lyumkis, Challenges and opportunities in cryo-EM single-particle analysis. J Biol Chem 294, 5181–5197 (2019).

15. D. Tegunov, L. Xue, C. Dienemann, P. Cramer, J. Mahamid, Multi-particle cryo-EM refinement with M visualizes ribosome-antibiotic complex at 3.5 Å in cells. Nature Methods 18, 186–193 (2021).

16. L. Jawerth et al., Protein condensates as aging Maxwell fluids. Science 370, 1317–1323 (2020).

17. M. Zhang et al., Molecular organization of the early stages of nucleosome phase separation visualized by cryo-electron tomography. Mol Cell 82, 3000–3014 e3009 (2022).

18. F. Tollervey et al., Cryo-Electron Tomography of Reconstituted Biomolecular Condensates. Methods Mol Biol 2563, 297–324 (2023).

19. M. Bose, M. Lampe, J. Mahamid, A. Ephrussi, Liquid-to-solid phase transition of oskar ribonucleoprotein granules is essential for their function in Drosophila embryonic development. Cell 185, 1308–1324.e1323 (2022).

20. X. Zhang et al., Molecular mechanisms of stress-induced reactivation in mumps virus condensates. Cell 186, 1877–1894.e1827 (2023).

21. H. Yu et al., HSP70 chaperones RNA-free TDP-43 into anisotropic intranuclear liquid spherical shells. Science 371, eabb4309 (2021).

22. F. J. B. Bauerlein et al., In Situ Architecture and Cellular Interactions of PolyQ Inclusions. Cell 171, 179–187 e110 (2017).

23. B. A. Gibson et al., Organization of Chromatin by Intrinsic and Regulated Phase Separation. Cell 179, 470–484.e421 (2019).

24. M. W. G. Schneider et al., A mitotic chromatin phase transition prevents perforation by microtubules. Nature (2022).

25. P. Li et al., Phase transitions in the assembly of multivalent signalling proteins. Nature 483, 336–340 (2012).

26. K. Luger, A. W. Mäder, R. K. Richmond, D. F. Sargent, T. J. Richmond, Crystal structure of the nucleosome core particle at 2.8 Å resolution. Nature 389, 251–260 (1997).

27. E. M. Hildebrand, J. Dekker, Mechanisms and Functions of Chromosome Compartmentalization. Trends Biochem Sci 45, 385–396 (2020).

28. A. M. Chiariello et al., Physical mechanisms of chromatin spatial organization. FEBS J 289, 1180–1190 (2022).

29. Y. Itoh, E. J. Woods, K. Minami, K. Maeshima, R. Collepardo-Guevara, Liquid-like chromatin in the cell: What can we learn from imaging and computational modeling? Current Opinion in Structural Biology 71, 123–135 (2021).

30. M. Eltsov et al., Nucleosome conformational variability in solution and in interphase nuclei evidenced by cryo-electron microscopy of vitreous sections. Nucleic Acids Res 46, 9189–9200 (2018).

31. F. Fatmaoui et al. (2022) Cryo-electron tomography and deep learning-based denoising reveal native chromatin landscapes of interphase nuclei. (Cell Biology).

32. Z. Hou, F. Nightingale, Y. Zhu, C. MacGregor-Chatwin, P. Zhang, Structure of native chromatin fibres revealed by Cryo-ET in situ. Nat Commun 14, 6324 (2023).

33. N. Jentink, C. Purnell, B. Kable, M. T. Swulius, S. A. Grigoryev, Cryoelectron tomography reveals the multiplex anatomy of condensed native chromatin and its unfolding by histone citrullination. Molecular Cell 83, 3236–3252.e3237 (2023).

34. Z. Y. Tan et al., Heterogeneous non-canonical nucleosomes predominate in yeast cells in situ. Elife 12 (2023).

35. A. J. Noble et al., Reducing effects of particle adsorption to the air– water interface in cryo-EM. Nature Methods 15, 793–795 (2018).

36. A. J. Noble et al., Routine single particle CryoEM sample and grid characterization by tomography. eLife 7 (2018).

37. K. Kelley et al., Waffle Method: A general and flexible approach for improving throughput in FIB-milling. Nature Communications 13, 1857 (2022).

38. K. Neselu et al., Measuring the effects of ice thickness on resolution in single particle cryo-EM. Journal of Structural Biology: X 7, 100085 (2023).

39. W. Wan, S. Khavnekar, J. Wagner, STOPGAP: an open-source package for template matching, subtomogram alignment and classification. Acta Crystallogr D Struct Biol 80, 336–349 (2024).

40. C. Purnell et al., Rapid Synthesis of Cryo-ET Data for Training Deep Learning Models. bioRxiv (2023).

41. T. Hrabe et al., PyTom: A python-based toolbox for localization of macromolecules in cryo-electron tomograms and subtomogram analysis. Journal of Structural Biology 178, 177–188 (2012).

42. D. Castano-Diez, M. Kudryashev, M. Arheit, H. Stahlberg, Dynamo: a flexible, user-friendly development tool for subtomogram averaging of cryo-EM data in high-performance computing environments. J Struct Biol 178, 139–151 (2012).

43. M. Beck et al., Visual proteomics of the human pathogen Leptospira interrogans. Nature Methods 6, 817–823 (2009).

44. I. Gubins et al. (2019) Classification in Cryo-Electron Tomograms. eds S. Biasotti, G. Lavoué, R. Veltkamp (The Eurographics Association).

45. S. Cruz-León et al., High-confidence 3D template matching for cryo-electron tomography. Nature Communications 15, 3992 (2024).

46. E. Moebel et al., Deep learning improves macromolecule identification in 3D cellular cryo-electron tomograms. Nature Methods 18, 1386–1394 (2021).

47. J. Singh, J. M. Thornton, The interaction between phenylalanine rings in proteins. FEBS Letters 191, 1–6 (1985).

48. D. Tegunov, P. Cramer, Real-time cryo-electron microscopy data preprocessing with Warp. Nature Methods 16, 1146–1152 (2019).

49. Y. T. Liu et al., Isotropic reconstruction for electron tomography with deep learning. Nat Commun 13, 6482 (2022).

50. D. Comaniciu, P. Meer, Mean shift: a robust approach toward feature space analysis. IEEE Transactions on Pattern Analysis and Machine Intelligence 24, 603–619 (2002).

51. J. Zivanov et al., A Bayesian approach to single-particle electron cryo-tomography in RELION-4.0. eLife 11, e83724 (2022).

52. R. V. Solé, S. Valverde, “Information Theory of Complex Networks: On Evolution and Architectural Constraints” in Complex Networks, E. Ben-Naim, H. Frauenfelder, Z. Toroczkai, Eds. (Springer Berlin Heidelberg, Berlin, Heidelberg, 2004), pp. 189–207.

53. F. Dar et al., Biomolecular condensates form spatially inhomogeneous network fluids. Nature Communications 15, 3413 (2024).

54. T. Wu et al., Single fluorogen imaging reveals distinct environmental and structural features of biomolecular condensates. bioRxiv, 2023.2001.2026.525727 (2024).

55. L. H. Wong, D. J. Tremethick, Multifunctional histone variants in genome function. Nat Rev Genet (2024).

56. G. Millan-Zambrano, A. Burton, A. J. Bannister, R. Schneider, Histone post-translational modifications - cause and consequence of genome function. Nat Rev Genet 23, 563–580 (2022).

57. N. J. Abdulhay et al., Massively multiplex single-molecule oligonucleosome footprinting. Elife 9 (2020).

58. S. Kale, A. Goncearenco, Y. Markov, D. Landsman, A. R. Panchenko, Molecular recognition of nucleosomes by binding partners. Curr Opin Struct Biol 56, 164–170 (2019).

59. M. Gelléri et al., True-to-scale DNA-density maps correlate with major accessibility differences between active and inactive chromatin. Cell Reports 42, 112567 (2023).

60. Anonymous, Chromatin arranges in chains of mesoscale domains with nanoscale functional topography independent of cohesin. SCIENCE ADVANCES, 17 (2020).

61. J. Bednar et al., Structure and Dynamics of a 197 bp Nucleosome in Complex with Linker Histone H1. Molecular Cell 66, 384–397.e388 (2017).

62. F. M. Richards, The interpretation of protein structures: Total volume, group volume distributions and packing density. Journal of Molecular Biology 82, 1–14 (1974).

63. T. Mittag, R. V. Pappu, A conceptual framework for understanding phase separation and addressing open questions and challenges. Mol Cell 82, 2201–2214 (2022).

64. N. Galvanetto et al., Mesoscale properties of biomolecular condensates emerging from protein chain dynamics. arXiv preprint arXiv:2407.19202 (2024).

65. Y. Shen et al., The liquid-to-solid transition of FUS is promoted by the condensate surface. Proceedings of the National Academy of Sciences 120, e2301366120 (2023).

66. M. Farag et al., Condensates formed by prion-like low-complexity domains have small-world network structures and interfaces defined by expanded conformations. Nature Communications 13, 7722 (2022).

67. Y. Dai et al., Biomolecular condensates regulate cellular electrochemical equilibria. Cell, S0092867424009097 (2024).

68. M. R. King et al., Macromolecular condensation organizes nucleolar sub-phases to set up a pH gradient. Cell 187, 1889–1906 e1824 (2024).

69. S. Ambadi Thody et al., Small-molecule properties define partitioning into biomolecular condensates. Nature Chemistry (2024).

70. C. Berger et al., Plasma FIB milling for the determination of structures in situ. Nature Communications 14, 629 (2023).

71. O. Schwartz et al., Laser phase plate for transmission electron microscopy. Nature Methods 16, 1016–1020 (2019).

72. Maria A. Ricci, C. Manzo, M. F. García-Parajo, M. Lakadamyali, Maria P. Cosma, Chromatin Fibers Are Formed by Heterogeneous Groups of Nucleosomes In Vivo. Cell 160, 1145–1158 (2015).

73. M. Lakadamyali, M. P. Cosma, Visualizing the genome in high resolution challenges our textbook understanding. Nature Methods 17, 371–379 (2020).

74. J. Xu et al., A comprehensive benchmarking with interpretation and operational guidance for the hierarchy of topologically associating domains. Nature Communications 15, 4376 (2024).

75. H. D. Ou et al., ChromEMT: Visualizing 3D chromatin structure and compaction in interphase and mitotic cells. Science 357, eaag0025 (2017).

76. O. H. Schiotz et al., Serial Lift-Out: sampling the molecular anatomy of whole organisms. Nat Methods 21, 1684–1692 (2024).

77. J. B. Woodruff et al., The Centrosome Is a Selective Condensate that Nucleates Microtubules by Concentrating Tubulin. Cell 169, 1066–1077.e1010 (2017).

78. B. R. Sabari et al., Coactivator condensation at super-enhancers links phase separation and gene control. Science 361, eaar3958 (2018).

79. J. Giudice, H. Jiang, Splicing regulation through biomolecular condensates and membraneless organelles. Nature Reviews Molecular Cell Biology 25, 683–700 (2024).

80. P. D. Dahlberg et al., Cryogenic single-molecule fluorescence annotations for electron tomography reveal in situ organization of key proteins in Caulobacter. Proceedings of the National Academy of Sciences 117, 13937–13944 (2020).

81. O. Klykov et al., In situ cryo-FIB/SEM Specimen Preparation Using the Waffle Method. Bio Protoc 12 (2022).

82. M. Schorb, I. Haberbosch, W. J. H. Hagen, Y. Schwab, D. N. Mastronarde, Software tools for automated transmission electron microscopy. Nat Methods 16, 471–477 (2019).

83. F. Eisenstein et al., Parallel cryo electron tomography on in situ lamellae. Nature Methods 20, 131–138 (2022).

84. S. Zheng et al., AreTomo: An integrated software package for automated marker-free, motion-corrected cryo-electron tomographic alignment and reconstruction. Journal of Structural Biology: X 6, 100068 (2022).

85. B. A. Lucas, N. Grigorieff, Quantification of gallium cryo-FIB milling damage in biological lamellae. Proc Natl Acad Sci U S A 120, e2301852120 (2023).

86. N. Sofroniew et al. (2024) napari: a multi-dimensional image viewer for Python. (Zenodo).

87. M. Chen et al., A complete data processing workflow for cryo-ET and subtomogram averaging. Nat Methods 16, 1161–1168 (2019).

88. D. N. Mastronarde, S. R. Held, Automated tilt series alignment and tomographic reconstruction in IMOD. Journal of Structural Biology 197, 102–113 (2017).

89. E. C. Meng et al., UCSF ChimeraX : Tools for structure building and analysis. Protein Science 32, e4792 (2023).

90. U. H. Ermel, S. M. Arghittu, A. S. Frangakis, ArtiaX : An electron tomography toolbox for the interactive handling of s ub-tomograms in UCSF ChimeraX. Protein Science 31 (2022).

91. A. A. Hagberg, D. A. Schult, P. J. Swart (2008) Exploring Network Structure, Dynamics, and Function using NetworkX. in Python in Science Conference, pp 11–15.

