## Supporting Information for "Quantitative Spatial Analysis of Chromatin Biomolecular Condensates using Cryo-Electron Tomography"

bioRxiv

**This PDF file includes:**

Figures S1 to S8
Tables S1

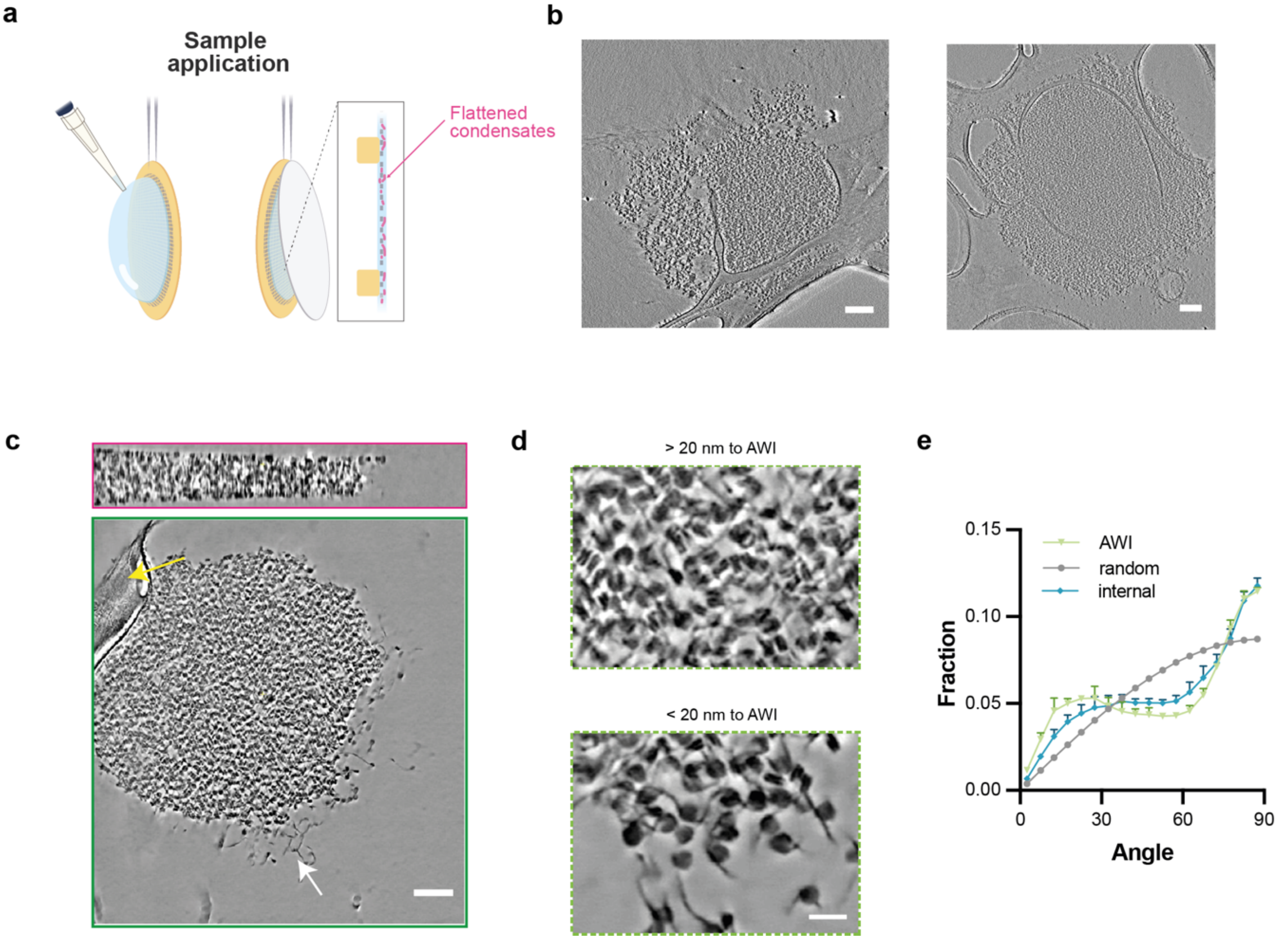


**Supplementary Figure 1. Disruption of condensate structures by back blotting and self-wicking technique**

**(a)** Diagram depicting the process of sample application followed by back blotting.

**(b)** A representative tomographic slice of a chromatin condensate prepared using the back blotting method. Scale bar is 100 nm.

**(c)** Orthogonal cross-sections of chromatin condensates processed using the self-wicking technique. Yellow arrows indicate carbon support film on the grids. White arrows indicate exposed nucleosomes and bare DNA regions. Scale bars are 100 nm. X-Y, X-Z views are shown with green, magenta outlines, respectively.

**(d)** Representative sections demonstrating chromatin condensates located close to and away from the air-water interface . Scale bars are 20 nm.

**(e)** Angular distribution relative to beam direction (Z-axis) of nucleosomes at the air-water interface and in the core of the condensate.

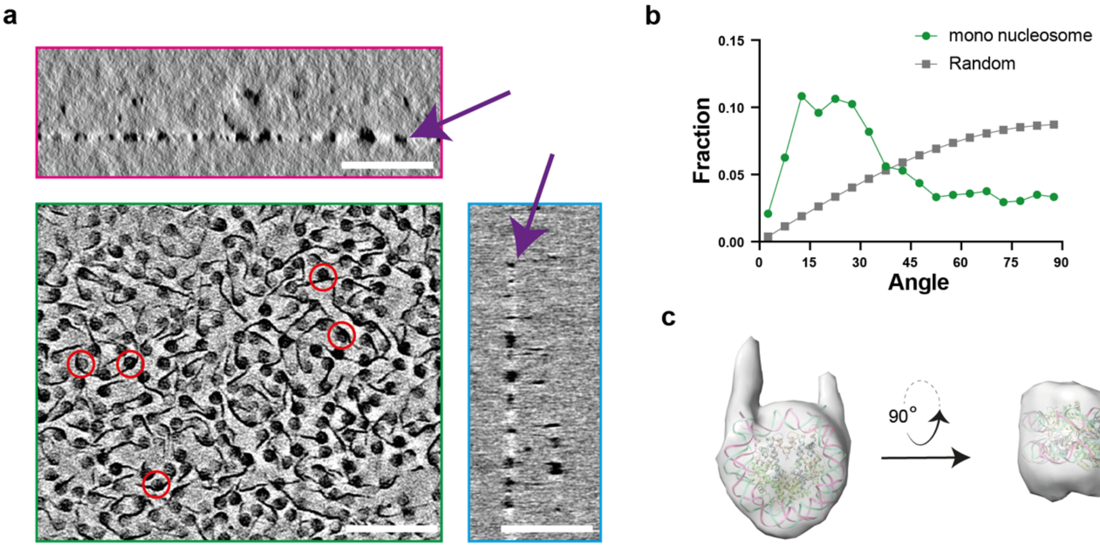


**Supplementary Figure 2. Denaturation and orientational bias of mononucleosomes at the air-water interface with blotting.**

**(a)** Orthogonal cross-sections from a reconstructed tomogram with mononucleosomes prepared with blotting. Purple arrows indicate the air-water interface. X-Y, X-Z and Y-Z views are shown with green, red and blue outlines, respectively. Red circles indicate some unwrapped nucleosomes. Scale bar is 100 nm.

**(b)** Distribution of nucleosome orientations with respect to the beam direction (Z-axis) as determined by CATM of the 4705 mononucleosomes observed in the tomograms.

**(c)** Subtomogram average structure of the mononucleosome from 4705 particles.

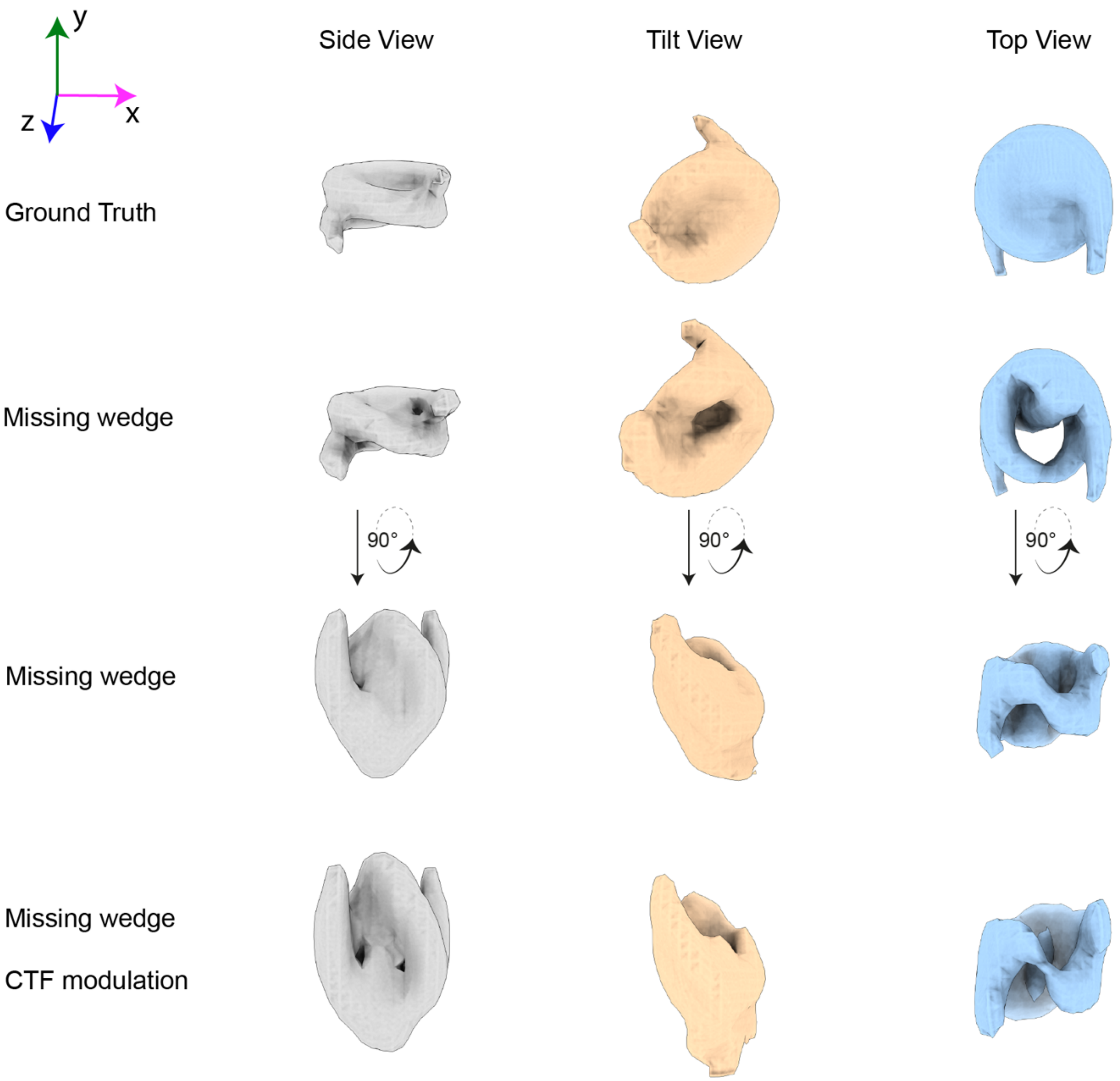


**Supplementary Figure 3. Missing wedge and CTF modulation distort the nucleosome**

The top panel displays a nucleosome structure from various perspectives. The missing wedge effect was then applied to nucleosomes in different orientations, causing pronounced distortions along the z-axis, as depicted in the bottom two panels. Additionally, the modulation by the contrast transfer function (CTF) further exacerbates the distortion, introducing more artifacts into the nucleosome images.

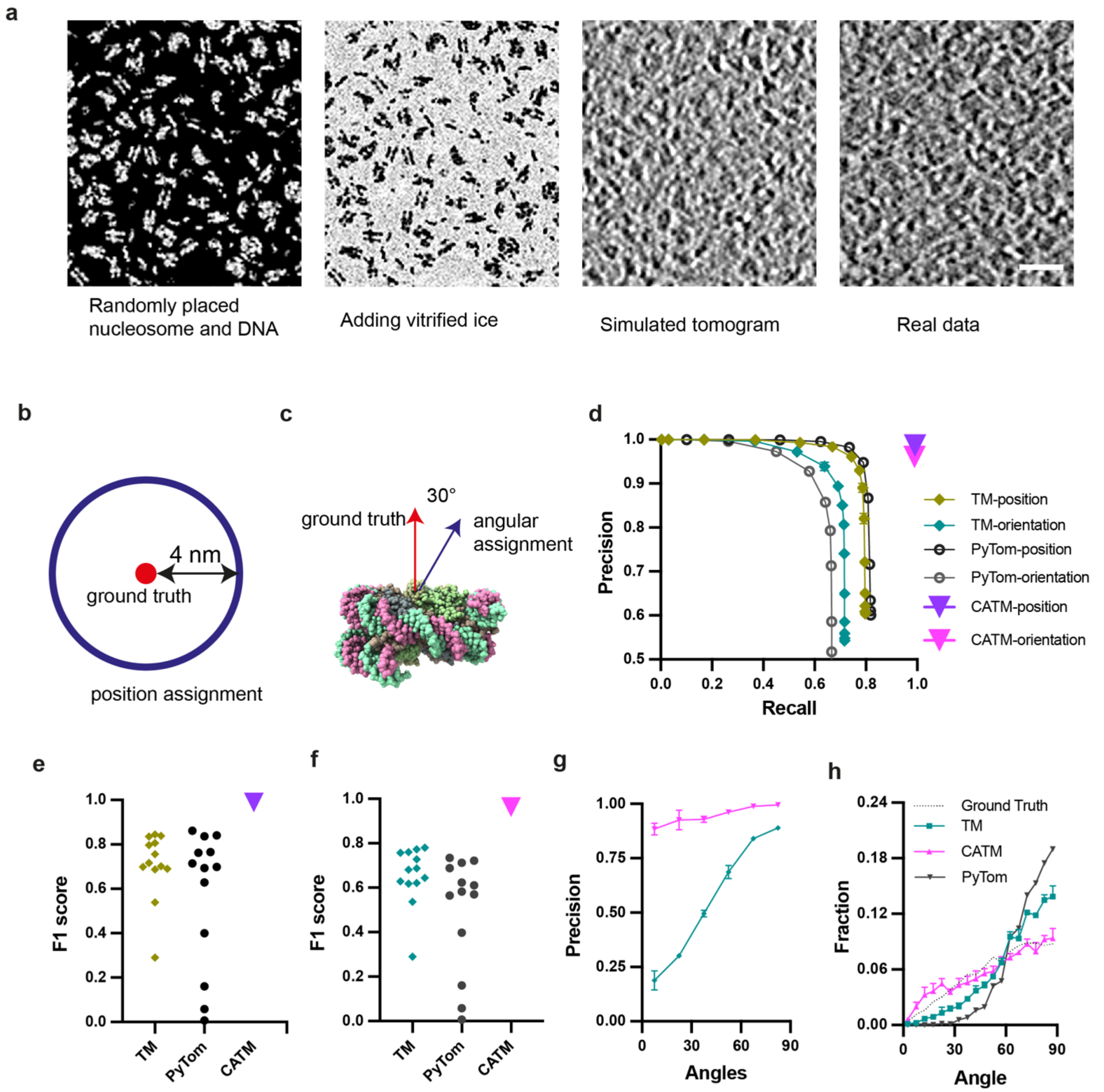


**Supplementary Figure 4. Benchmark of CATM against standard template matching and PyTom template matching.**
**(a)** Simulation of tomograms. The nucleosomes and 25 bp DNA were randomly placed in a 3D volume at a density equivalent to chromatin condensates, and vitrified ice was simulated. The volume was then subjected to tilt-series simulation and tomogram reconstruction. The tomograms was low pass filtered to 25 Å and compared with real chromatin data with the same filter. Scale bar is 20 nm.
**(b)** Particles with center of mass assigned within 4 nm of the center of mass of a ground truth nucleosome are considered accurate.
**(c)** For orientation accuracy, a vector perpendicular to the nucleosome plane was defined. The angle between the vectors from the ground truth and assigned nucleosomes was calculated. Predictions within 30° were considered accurate.
**(d)** Comparative performance in nucleosome assignment by CATM, TM and PyTom. For TM and PyTom, with various of cross-correlation cutoffs the algorithms have different performance. CATM produces only a single set of assignments.
**(e-f)** F1 scores of position (e) and orientation (f) for the different algorithms.
**(g)** Orientation assignment precision as a function of orientation with respect to the beam direction (Z-axis) for each method. Zero degrees is the top view of a nucleosome (disc oriented perpendicular to beam direction), 90 degrees is a side view.
**(h)** Distribution of nucleosome orientations with respect to the beam direction (Z-axis) as determined by CATM, TM and PyTom. Nucleosomes are oriented randomly in the ground truth (dash line), producing a sinusoidal distribution.

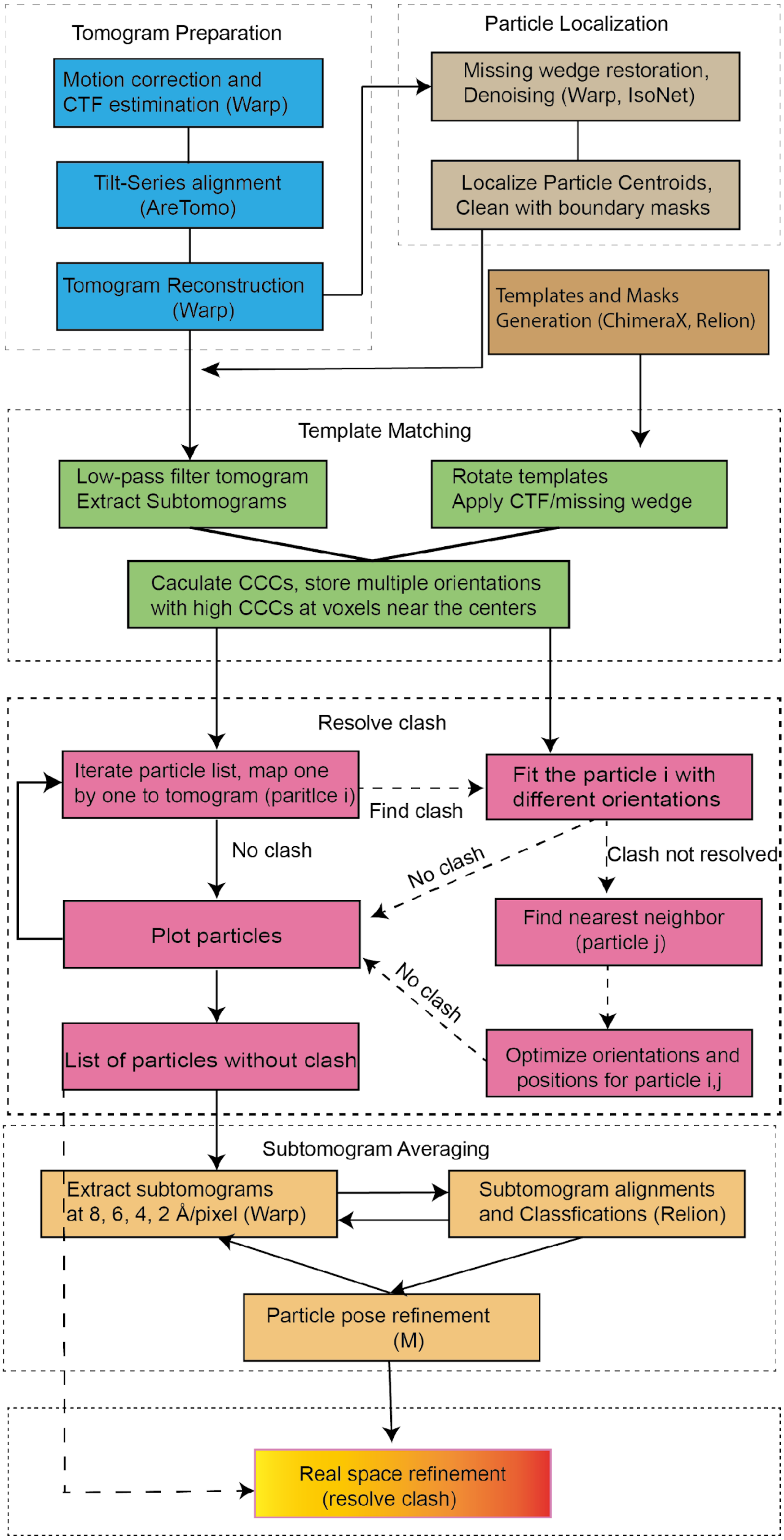


**Supplementary Figure 5. CATM data analysis pipeline**

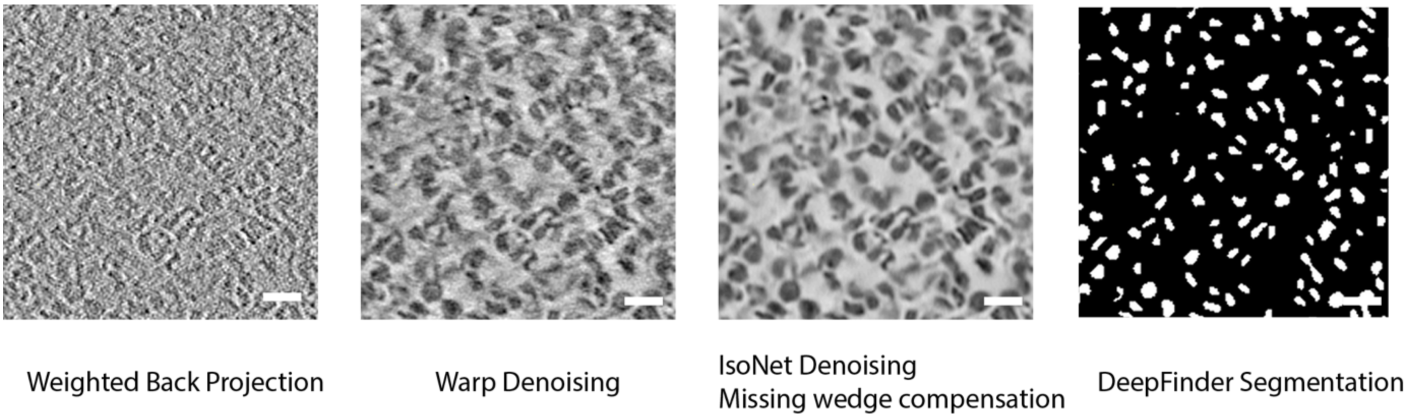


**Supplementary Figure 6. Tomogram denoising, missing wedge compensation, and segmentation.**
Representative slices of tomogram generated by weighted back projection, Warp and IsoNet denoising , missing wedge compensation, and the DeepFinder segmentation. Scale bar is 20 nm.

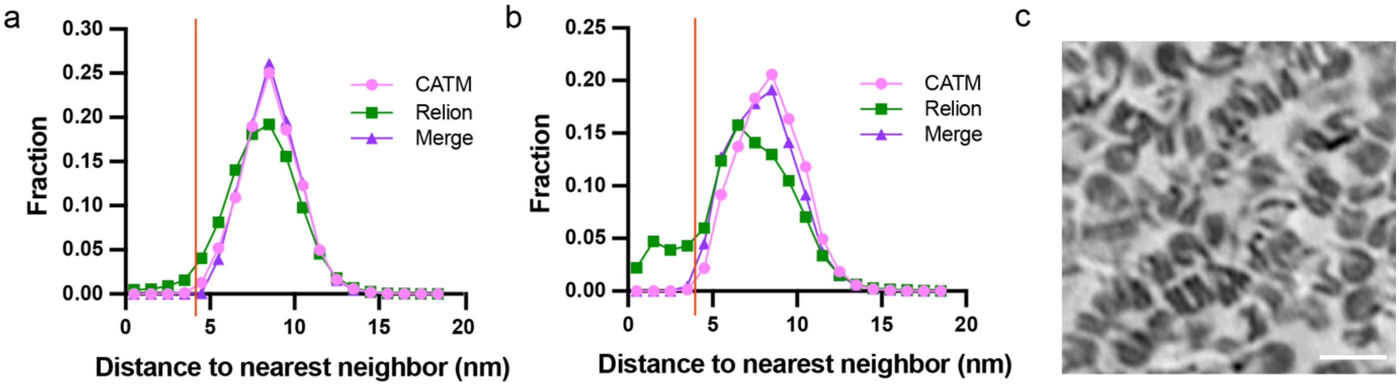


**Supplementary Figure 7. Relion tends to refine particles into local minima**

Quantification of the nearest neighbor distance for CATM, Relion and the subsequent refined particles which merge the CATM and Relion refinement for chromatin condensate **(a)** and another case of chromatin condensates enriched in stacking nucleosomes **(b).** The yellow vertical line indicates the nucleosome distance tolerance of 4 nm, corresponding to the width of the nucleosome disk.

**(c)** A representative image of the condensate with highly abundant stacking.

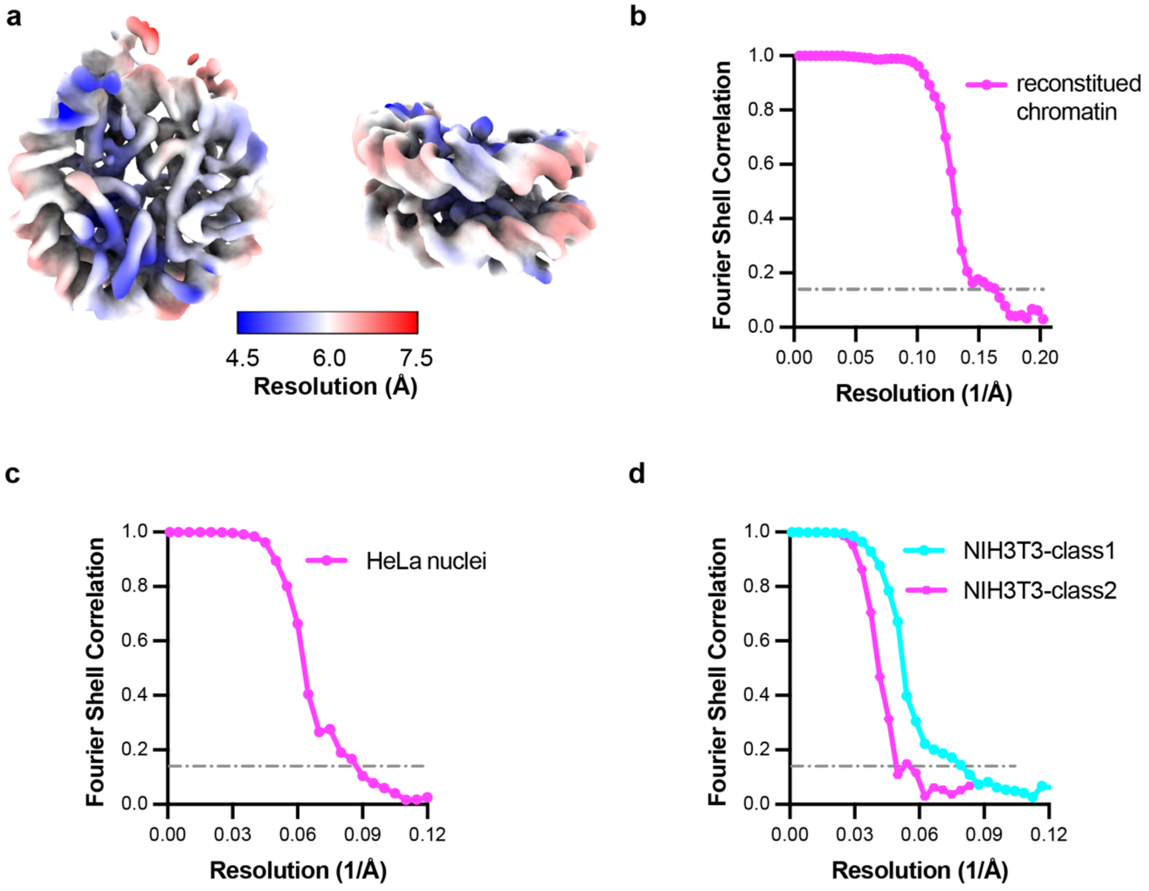


**Supplementary Figure 8. Resolution estimation for subtomogram averaging with FSC**
**(a)** Subtomogram averaging to reconstruct a nucleosome structure from reconstituted chromatin condensates, colored by local resolution.
**(b-d)** Fourier shell correlation between two independently processed halves of the data set used in reconstituted chromatin condensates (b), HeLa nuclei (c) and NIH3T3 (d) tomograms.

**Supplementary Table 1: cryo-ET data collection and reconstruction statistics**

| | Vitrobot-prepared chromatin | Chameleon prepared | HPF prepared Reconstituted chromatin | Nuclei chromatin | Cellular chromatin |
|---|---|---|---|---|---|
| Data collection | | | | | |
| Magnification | 33 K | 33 K | 81 K | 81 K | 64 K |
| Voltage (kV) | 300 | 300 | 300 | 300 | 300 |
| Defocus range (µm) | -0.5 | -0.5 | -3 to -4.5 | -3 to -4.5 | -4 |
| Pixel size (Å) | 2.06 | 2.06 | 1.516 | 1.516 | 1.516 |
| Electron exposure (e–/Å$^2$) | 150 | 150 | 178 | 178 | 178 |
| Tilt-range/step (°) | -60,+60 / 2 | -60,+60 / 2 | -48,+60 / 2 | -48,+60 / 2 | -54,+54 / 3 |
| Tilt-scheme | dose-symmetric, grouping 2 | | dose-symmetric, grouping 3 | | dose-symmetric, grouping 1 |
| Processing | | | | | |
| Symmetry imposed | - | - | C1 | C1 | C1 |
| Initial particle images (no.) | - | - | 126125 | 35503 | 13277 |
| Final particle images (no.) | - | - | 126125 | 35503 | Class 1: 6470<br>Class 2: 6804 |
| Map resolution (Å) | - | - | 6.1 | 12 | Class1: 12<br>Class2: 22 |
| FSC threshold | 0.143 | | | | |
